# Characterizing dynamic tissue architectures by identifying cell-type-specific spatiotemporal gene programs with stGP

**DOI:** 10.64898/2026.07.03.736035

**Authors:** Baichen Yu, Ziyue Tan, Xiaomeng Wan, Hansheng Wang, Can Yang

## Abstract

Cellular gene programs unfold over biological time within spatially organized tissues. The same cell type can activate distinct programs in different spatial domains or multicellular niches. Spatiotemporal transcriptomics enables in situ measurement of these processes; however, the interplay between temporal progression and spatial organization complicates the identification of whether a gene program is influenced by temporal changes, spatial structure, or both. To overcome this challenge, we present spatiotemporal Gene Programs (stGP), a statistical framework for identifying interpretable cell-type-specific gene programs across multi-sample spatiotemporal transcriptomic studies. stGP preserves the molecular identity of each program through shared gene loadings, and decomposes individual cell activity into a temporal component that captures gene program responses over biological time, and a spatial component that characterizes variations within tissue sections. We quantify their relative contributions by estimating their variance components. Through comprehensive simulations and analyses of three spatiotemporal transcriptomic datasets across different tissues and technologies, stGP reveals spatiotemporal gene programs that distinguish preserved tissue architecture from dynamic remodeling. Our framework delineates age-associated cellular responses and uncovers localized program deployment within anatomical regions, aging hotspots, and multicellular niches. Our results establish stGP as an effective and robust framework for dissecting dynamic tissue architecture, providing insights into how cell-type-specific gene programs are coordinated across biological time and spatial microenvironments.

## Introduction

Cellular gene programs evolve over biological time within spatially organized tissues^1-3^. During biological processes such as aging^4,5^, development^6^, injury and repair^7,8^, the same cell type can activate different molecular programs at different biological stages, in distinct anatomical regions or within specific multicellular neighborhoods. Recent spatiotemporal transcriptomic atlases enable the in situ measurement of these processes, providing an unprecedented opportunity to link gene programs to cell identity, tissue architecture and biological time^9-11^. However, the analysis of these datasets presents a significant challenge: temporal progression at the sample level, anatomical organization within each tissue section, and local niche-associated variation occur simultaneously across distinct and often unaligned tissue sections. Thus, the key question arises: does a gene program change with age or stage, and does its activity reflect a temporal response shared across samples, a stable spatial domain, or the localized deployment of a common temporal process? To address this question, it requires a model that treats time and space as structured but separable sources of variation, while preserving the molecular identity of each gene program.

Currently, most existing approaches only partially address this need. Classical factorization methods, including PCA and NMF^12^, can capture variation with interpretable embeddings, but they do not distinguish whether a latent factor is driven by spatial organization, biological time or their interaction. Spatially aware methods such as SpatialPCA^13^, SpiceMix^14^, and Popari^15^ incorporate spatial structure to improve tissue-level representation or multi-sample spatial factorization, and STAMP extends topic modeling to time-series spatial transcriptomic data^16^. MEFISTO models smooth variation along temporal or spatial covariates in multi-view data^17^. Nevertheless, most of these methods lack a unified framework for decomposing cell-type-specific program activity into temporal trajectories and spatial effects while identifying interpretable gene loadings. As a result, they fall short in addressing key questions central to dynamic tissue biology, such as how a program changes across age, how strongly it is organized in space, and whether its activation reflects broad temporal drift or anatomically localized remodeling.

To overcome the challenge, we here developed spatiotemporal Gene Programs (stGP), a statistical framework for identifying interpretable cell-type-specific gene programs in multi-sample spatiotemporal transcriptomic studies. stGP is built on the principle that a gene program can retain a stable molecular identity across samples, whereas its activity may vary with biological time and local spatial context. For a target cell type, stGP represents gene expression as a small set of latent programs with non-negative gene loadings on a simplex, making each program interpretable as a weighted gene set shared across samples. For each program, stGP decomposes per-cell activity into a temporal trajectory shared by cells from samples at the same biological age, and a within-section spatial effect that captures local program deployment across tissue coordinates. Gaussian process priors are used to smooth these components over biological time and within-section spatial coordinates, while variance components quantify their relative contributions.

We evaluated stGP through comprehensive simulations and three spatiotemporal transcriptomic studies spanning different tissues, biological processes and technologies (e.g., MERFISH^18^ and Xenium^19^). Across these settings, stGP showed that cell-type-specific programs can follow coherent temporal trajectories across samples while being selectively deployed within anatomical domains, aging-associated hotspots or multicellular niches. By preserving shared gene loadings and decomposing program activity into temporal and within-section spatial components, stGP separates age-associated responses from local tissue organization and links each program to interpretable molecular signatures. In human dorsolateral prefrontal cortex sections spanning a broad age range, stGP resolved laminar excitatory-neuron architecture more accurately than baseline methods and identified an oligodendrocyte program associated with age-dependent multicellular niches. In aging mouse brains, stGP uncovered a late-life microglial MHC-I program enriched in white-matter regions with T-cell proximity effects. In a bilateral mouse kidney injury-repair time course, stGP identified replicable and robust repairing programs across biological replicates. Together, these results support stGP as an effective and interpretable framework for converting spatiotemporal transcriptomic data into temporal trajectories and spatially resolved models of dynamic tissue architecture, enabling systematic investigation of how cell-type-specific molecular programs are coordinated across biological time, anatomical space, and local tissue microenvironments.

## Results

### Overview of stGP

stGP is a statistical framework designed for multi-sample spatiotemporal transcriptomic studies, where cells are observed across a temporal covariate such as chronological age and within two-dimensional tissue spatial coordinates (Fig. 1). The model is fitted separately within each annotated cell type. For a target cell type, stGP takes a cell-by-gene expression matrix for each sample, the sample-level temporal covariate and the within-section spatial coordinates of each cell as input (Fig. 1a). It represents expression data as a small number of latent gene programs, each defined by a non-negative sum-one loading vector (Fig. 1a and Methods). This makes each program interpretable as a weighted gene set rather than an arbitrary signed factor.

**Fig. 1.**
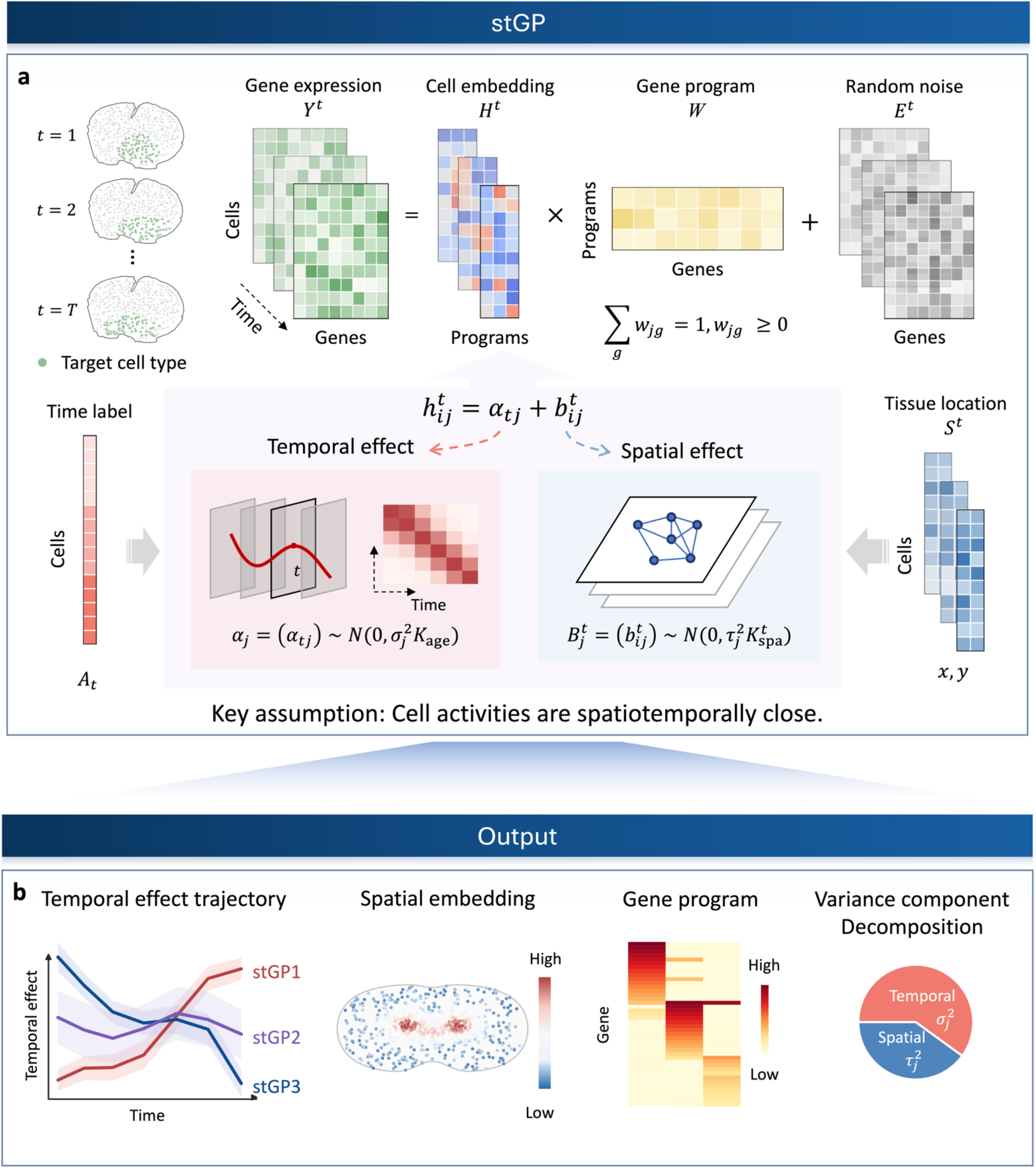
Overview of stGP. a, The statistical model of stGP. stGP is designed for multi-sample spatiotemporal transcriptomic studies measured across biological time and is fitted within a target cell type. stGP represents expression data as a small number of latent gene programs. The program-specific cell activity is decomposed into a sample-level temporal effect and a within-section spatial effect, with Gaussian process priors. **b**, The output of stGP. stGP can decompose temporal effects and heterogeneous spatial embeddings, and produce both interpretable gene program loadings and variance component decomposition quantifying the relative temporal and spatial contributions.

For each gene program, the activity of each cell is decomposed into two components (Fig. 1a). The first is a sample-level temporal effect shared by all cells of the same cell type from a given individual. This component is smoothed across the temporal covariate by a Gaussian process prior, which describes the synchronizing temporal effect shared by cells from samples with the same chronological age or stage^20^. The second is a cell-level heterogeneous spatial effect that varies across locations within each tissue section. This component is smoothed by another spatial Gaussian process prior defined on pairwise distances within the section (Methods). The key idea of this decomposition is to assume that cell activities in each gene program are spatiotemporally close, i.e., cells with similar biological age and within-slice spatial position are likely to have similar latent embeddings. This construction separates variation across individuals from anatomical variation within each section and avoids the need to register sections to a common coordinate system. The identifiability issue for stGP is addressed since the priors are well placed and the weight matrix is nonnegative (Supplementary Methods A.5).

stGP estimates gene loadings, cell embeddings and variance components using a blockwise backfitting algorithm (Methods and Supplementary Methods A). For each rank-1 update, the algorithm alternates between updating program activities, gene loadings on the simplex, and variance components. Multi-program models are obtained by sequentially extracting rank-1 components from residuals, with negligible or near-duplicate programs removed during model selection. Notably, the number of gene programs could be automatically determined in the blockwise backfitting algorithm (Supplementary Methods A.6).

For each program, the fitted model returns a gene-loading vector, a temporal trajectory with posterior uncertainty, section-specific spatial embeddings and variance component estimates (Fig. 1b). These outputs support direct biological interpretation. The variance fractions quantify the relative contributions of age and space for every gene program. High-weight genes define the molecular identity of a program, temporal trajectories reveal when the program is activated or repressed, and spatial embeddings show where the program localizes within each tissue section. Downstream analyses use these outputs to annotate gene programs, compare spatial organization across ages or donors, identify temporally or spatially dominated programs, cluster the spatial domains, and relate program activity to local cell neighborhoods or multicellular niches.

### stGP accurately recovers spatiotemporal gene programs in simulation studies

We conducted comprehensive simulation experiments to benchmark stGP against baseline methods for spatiotemporal gene program modeling (Fig. 2). The compared methods in the simulation study include gene module detection methods either considering spatial or spatiotemporal dependence (SpatialPCA^13^, Popari^15^, STAMP^16^, and MEFISTO^17^) and two dimension reduction methods (PCA and NMF^12^). For all methods, we set the number of gene programs as the true value p = 4. The automatic selection of the number of programs can be done by the stGP backfitting algorithm (Supplementary Fig. S1). Among these four programs, we first simulate the signals on the log-scale, two of them are mixed with both spatial and temporal signals, one is purely spatial, and one is purely temporal. Then we used Poisson distribution to generate count data to mimic the spatial transcriptomic counts. stGP also supports accurate recovery results in the direct log-scale fitting (Supplementary Fig. S3). The details of simulation designs are provided in Supplementary Notes B.2.

**Fig. 2.**
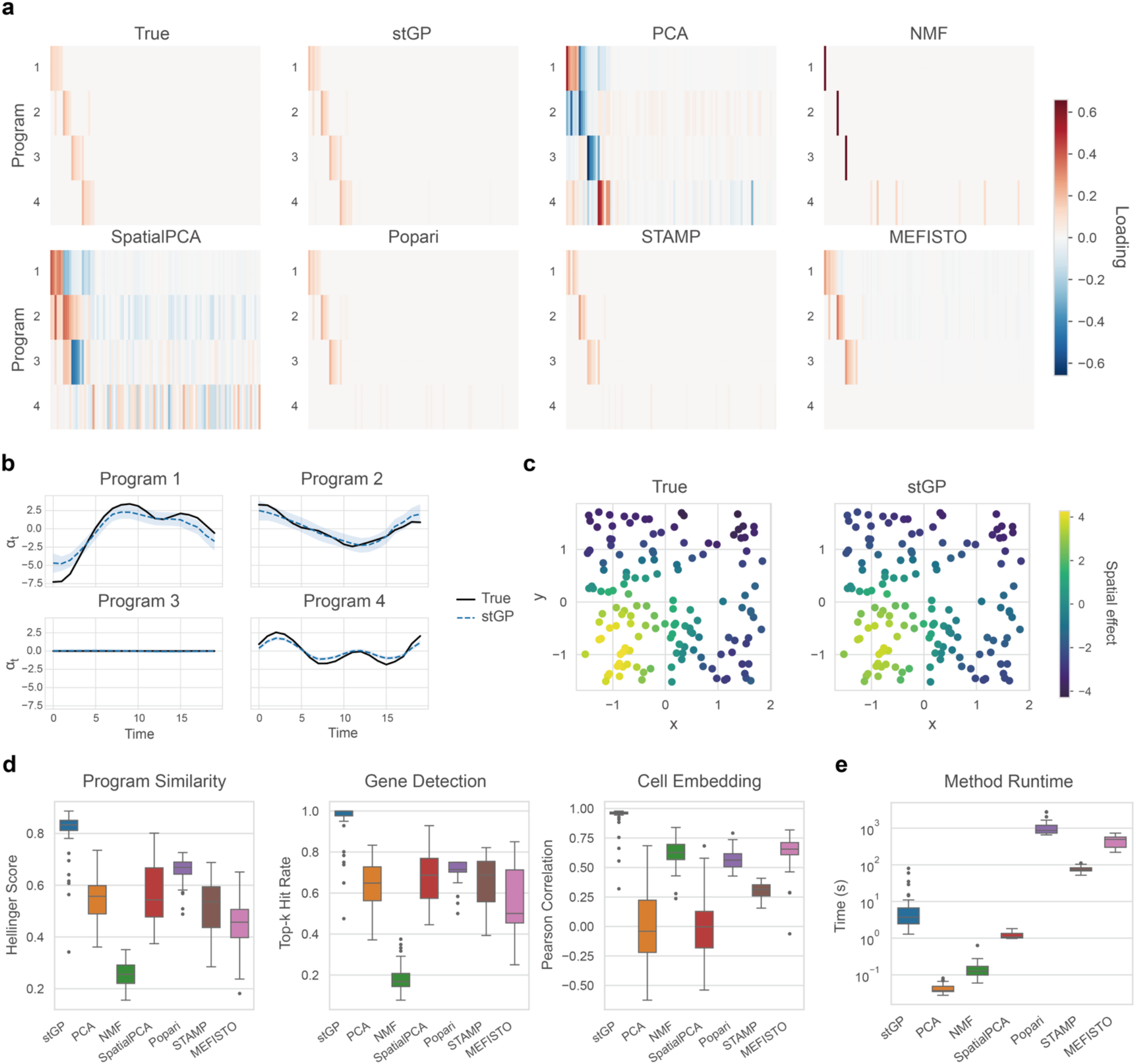
stGP recovers spatiotemporal gene programs by disentangling spatial and temporal effects in simulated data. **a**, True and estimated gene loadings for stGP, PCA, NMF, SpatialPCA, Popari, STAMP and MEFISTO in one representative replicate. **b**, Recovered temporal effects for the four latent programs of stGP, with 95% posterior intervals for uncertainty quantification. **c**, True and stGP-recovered within-section activity maps in spatial coordinates. **d**, Quantitative metrics comparing stGP with baseline methods across 50 simulation replicates. The program similarity is measured by Hellinger score, gene detection ability is measured by the top-k hit rate, and cell embeddings are evaluated by Pearson correlation, respectively. **e**, Runtime comparison across methods.

stGP recovered the latent gene programs more accurately than the baseline methods. In a representative replicate, the simplex-constrained loadings learned by stGP closely matched the ground-truth programs and preserved clear separation among the four programs (Fig. 2a). In contrast, PCA and SpatialPCA produced mixed loadings due to their orthogonality constraints. NMF recovered only one of the most variable genes in each program, whereas Popari, STAMP and MEFISTO showed weaker recovery of programs dominated by sample-level temporal variation. Particularly, none of the existing methods can recover the gene program with only temporal effects (Fig. 2a). Based on 50 replications under this simulation setting, stGP achieved the best quantitative results in terms of gene program identification, gene detection, and cell embedding recovery (Fig. 2d). Additional evaluation metrics supported the same conclusion (Supplementary Fig. S5).

The main advantage of stGP was that it recovered temporal and spatial components as separate, interpretable outputs after removing the temporal confounders. stGP reconstructed the temporal trajectories of latent programs with uncertainty quantification and correctly identified programs with temporal effects, spatial effects, or both (Fig. 2b). It also recovered the within-section spatial activity in the tissue coordinates, highlighting the activated bottom left region of one particular gene program (Fig. 2c).

stGP also remained practical and robust under realistic modeling choices. Due to efficient computational techniques (Methods and Supplementary Methods A.2.2), stGP was clearly faster than Popari^15^, STAMP^16^, and MEFISTO^17^ (Fig. 2e). We also showed the robustness of stGP against various kernel misspecifications (Supplementary Fig. S2), and under the single-cell-like settings without spatial effects (Supplementary Fig. S4). Together, these simulations show that stGP recovers interpretable gene programs while deciphering temporal and spatial variations across different settings.

### stGP outperforms existing methods in resolving laminar architecture in unaligned human DLPFC sections

We initially considered a MERFISH dataset from a human dorsolateral prefrontal cortex (DLPFC), which included 12 sections from donors aged 15–87 years^5^. Here we focused on excitatory neurons, which are predominantly glutamatergic projection neurons and exhibit strong laminar specialization across cortical layers^21^. This cell population therefore provides a biologically grounded benchmark for evaluating whether stGP can recover layer-resolved spatial organization from unaligned tissue sections.

We applied stGP to the DLPFC dataset, and identified four layer-specific gene programs in excitatory neurons with distinct regional activities and spatiotemporal strength (Fig. 3f and Supplementary Fig. S6). Three programs correspond to three major laminar layers of the cortex, respectively (Fig. 3d and Supplementary Fig. S7-S9). Specifically, stGP1 marked the upper L2/3 layer and was enriched for the canonical marker *CUX2* (Supplementary Fig. S7), stGP3 marked middle L4 layer with *RORB* as the marker gene (Fig. 3b,d and Supplementary Fig. S8), and stGP2 marked the deep L5/6 layer and the gray-white matter boundary of DLPFC, with *HS3ST4* as a representative marker gene (Supplementary Fig. S9)^5^. Clustering analysis based on the stGP spatial embedding also produced spatial patterns similar to the layer structure (Fig. 3a-c). This not only shows that stGP is able to recover the spatial tissue architecture, but also reveals that the cortex layer structure is rather stable across lifespan. Moreover, increasing the number of clusters using the same stGP embedding further resolved finer sublaminar domains within the major layers (Supplementary Fig. S10), indicating that stGP retains additional spatial information beyond a coarse three-layer partition. The temporal effect of stGP3 showed a modest decreasing trend with age (Fig. 3i), suggesting that the laminar identity remained spatially preserved whereas the corresponding transcriptional state was attenuated with aging. The deep-layer program stGP2 showed a non-monotonic age trajectory, with higher activity in midlife and reduced activity in older donors (Supplementary Fig. S6). This pattern similarly suggests that deep-layer excitatory neurons preserve their anatomical position while their layer-associated transcriptional program changes with aging.

**Fig. 3.**
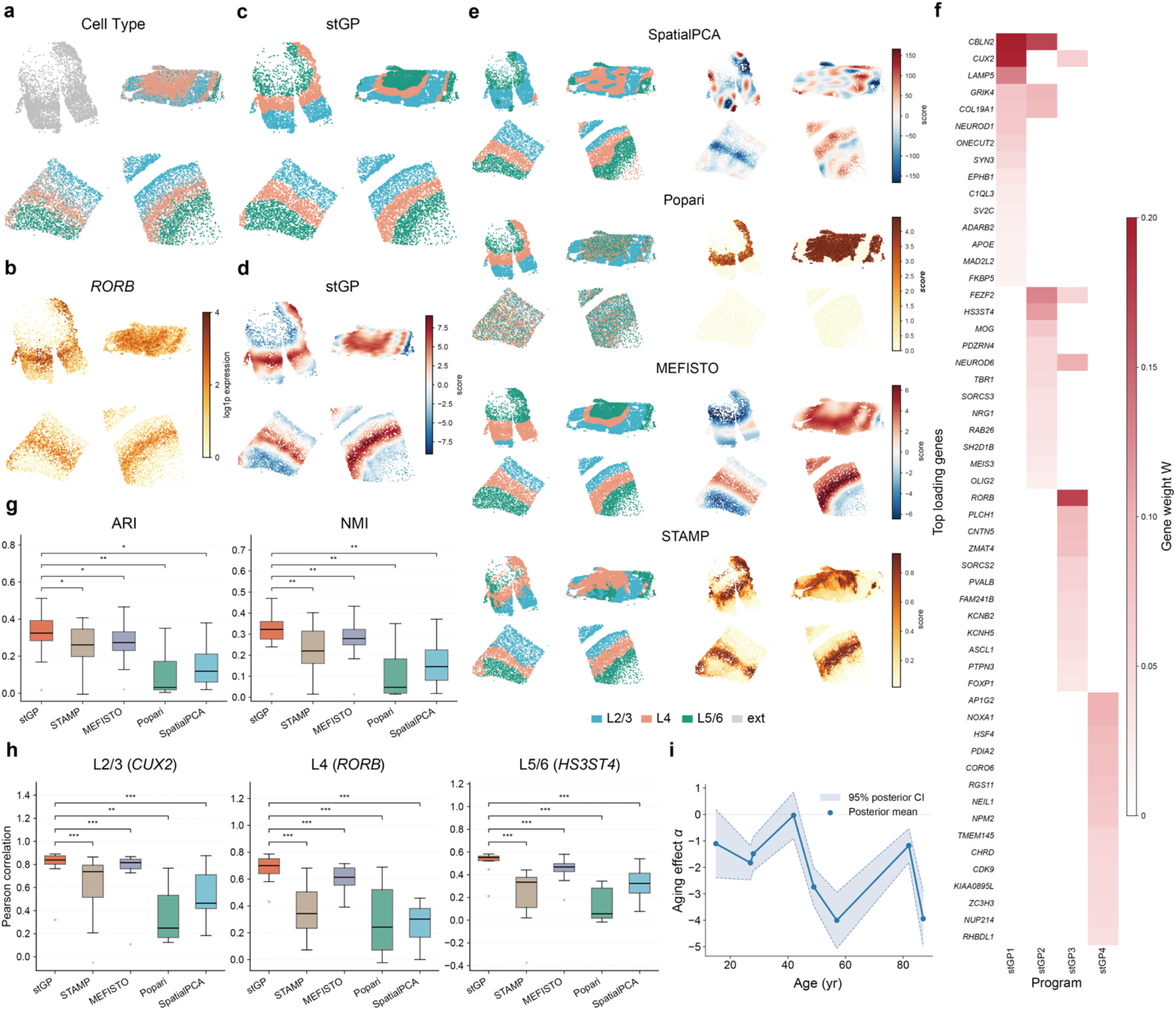
stGP improves recovery of laminar excitatory-neuron architecture in unaligned aging human DLPFC MERFISH sections. **a**, Reference excitatory-neuron cell subtype annotations for representative DLPFC sections with different ages. Grey denotes cells without a layer-specific subtype assignment. **b**, Spatial log1p-transformed expression of the L4-layer marker *RORB*. **c**, The laminar domains inferred by clustering stGP spatial embeddings into three domains. **d**, The L4-layer associated stGP3 program spatial activities, showing concordant localization to the L4 band. **e**, Spatial domains (left panels) inferred by baseline methods and the method-specific latent scores most associated with the L4 layer (right panels). From top to bottom: SpatialPCA, Popari, MEFISTO and STAMP. **f**, stGP gene program loading matrix across four programs. **g**, Quantitative comparison of layer recovery using ARI (left panel) and NMI (right panel) against available marker-defined domains across sections. **h**, Pearson correlations between method embeddings and layer marker expressions for L2/3 (*CUX2*), L4 (*RORB*) and L5/6 (*HS3ST4*). Statistical significance was assessed at the section level using one-sided paired Wilcoxon signed-rank tests (*, *P* < 0.05; **, *P* < 0.01; ***, *P* < 0.001). **i**, Estimated temporal effect for the stGP3 L4-layer associated program across donor age.

To benchmark the ability of resolving spatial patterns in multi-sample spatiotemporal transcriptomic studies, we compared stGP with SpatialPCA^13^, MEFISTO^17^, Popari^15^, and STAMP^16^ (Methods and Supplementary Notes B.1). We assessed continuous embeddings by Pearson correlation with matched layer-marker expression, including *CUX2* for L2/3, *RORB* for L4 and *HS3ST4* for L5/6^5^. We further evaluated clustered spatial domains using adjusted Rand index (ARI)^22^ and normalized mutual information (NMI)^23^ against marker-defined layers and, where available, cell subtype labels of excitatory neurons (Methods). By separating temporal effects from spatial organization, we found that stGP consistently outperformed all baseline methods across all metrics (Fig. 3). Specifically, stGP captured both broad laminar bands and fine-scale boundary details across sections with different ages (Fig. 3c,d). SpatialPCA produced fragmented spatial components and patchy clusters, due to the absence of temporal modeling (Fig. 3e and Supplementary Fig. S7-S9). Although Popari is designed for multisample spatial decomposition, it recovered laminar structure only in a few sections and missed clear layer organization in others (Fig. 3e and Supplementary Fig. S7-S9). STAMP and MEFISTO recovered some broad layer-like gradients but showed reduced continuity or boundary precision in several sections (Fig. 3e and Supplementary Fig. S7-S9). Quantitative results further supported the improved performance of stGP (Fig. 3g,h). More specifically, stGP achieved the highest marker correlations across all layer markers (Fig. 3h), as well as the highest ARI and NMI for spatial domains defined by marker genes and available subtype annotations (Fig. 3g and Supplementary Fig. S10). Thus, in unaligned human DLPFC sections spanning a broad age range, stGP more accurately resolves layer-specific excitatory-neuron architecture while revealing interpretable age-dependent activity for each program.

### stGP uncovers the aging effect of multicellular niches in human brain oligodendrocytes

We further explored the DLPFC analysis results from stGP to uncover aging effects on multicellular niches in the human brain. For oligodendrocytes, stGP identified a spatially restricted program, stGP4, with a pronounced age-associated temporal component (Fig. 4c,d). Because oligodendrocytes function within glial-rich tissue compartments rather than in isolation, we next asked whether stGP4 reflected a purely cell-intrinsic oligodendrocyte state or a program coupled to the local multicellular microenvironment. To address this issue, we applied NicheScope^24^, which characterizes shared and young- or aged-specific multicellular niches (MCNs) and their corresponding niche-regulated cell states from spatial transcriptomics data, using oligodendrocytes as the target cell type.

**Fig. 4.**
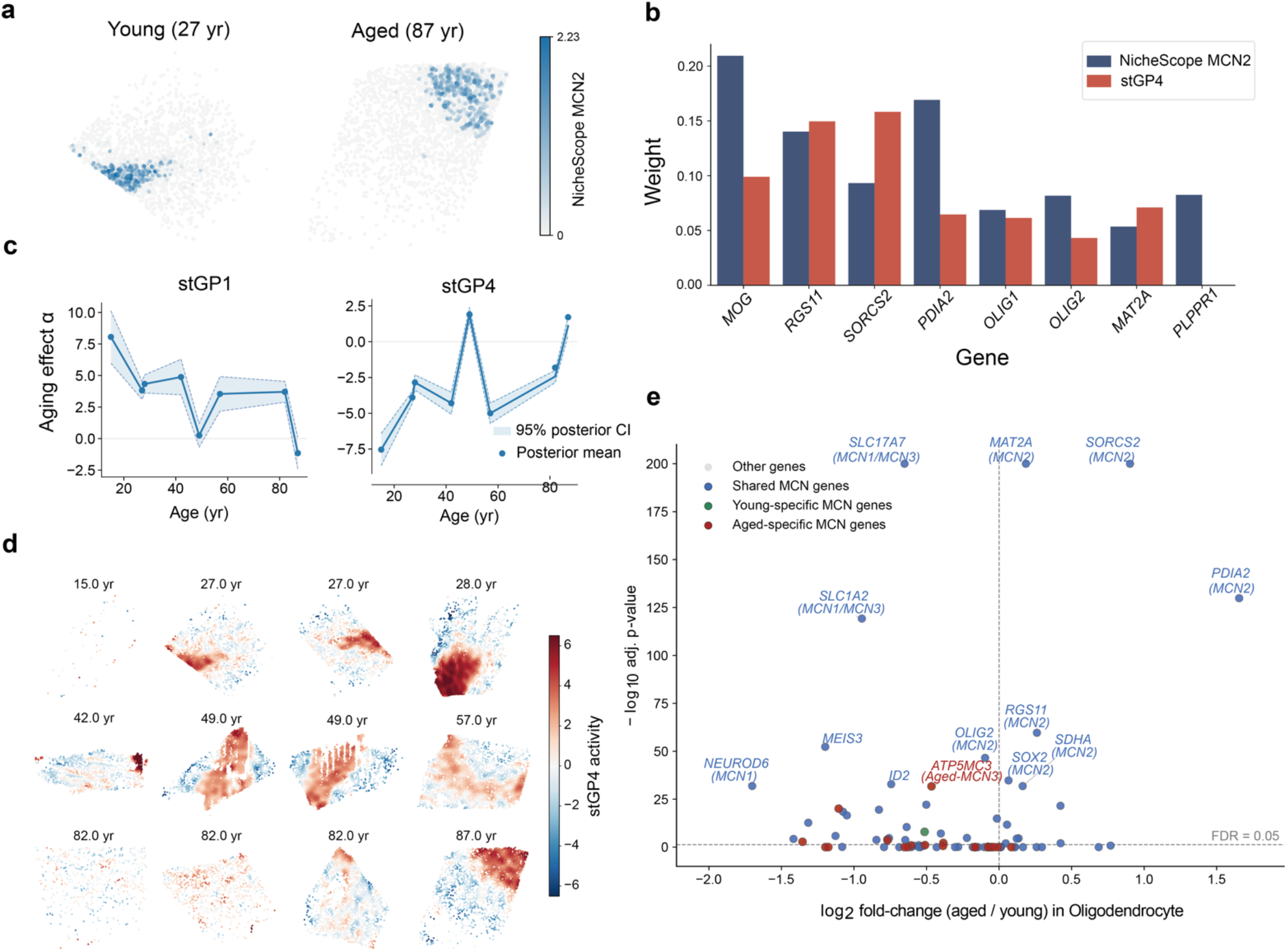
stGP reveals age-associated remodeling of a shared oligodendrocyte multicellular niche in human DLPFC. **a**, Spatial distribution of the NicheScope MCN2 niche score in representative young and aged DLPFC sections. **b**, Comparison of top gene weights between the NicheScope MCN2 niche-regulated state and stGP4. **c**, Estimated age-associated temporal effect of stGP1 (left panel) and stGP4 (right panel) across donors. **d**, Spatial maps of stGP4 scores across DLPFC sections ordered by donor age. **e**, Differential expression analysis comparing young versus aged oligodendrocytes. Genes are colored according to their classification as shared MCN genes, young-specific MCN genes, aged-specific MCN genes or other genes. The vertical dashed line marks a log2 fold-change of zero, and the horizontal dashed line marks FDR = 0.05. Labels indicate representative highly significant MCN-associated genes.

Among the niches inferred by NicheScope, shared MCN2 showed the strongest correspondence with stGP4 (Fig. 4b). Top-weighted genes from the NicheScope MCN2 niche-regulated state overlapped with high-loading stGP4 genes, including oligodendrocyte and myelin-associated genes such as *MOG, OLIG1* and *OLIG2*, together with *PDIA2* and *RGS11* (Fig. 4b and Supplementary Figs. S11,S12). This concordance indicates that stGP4 captures the molecular axis of an oligodendrocyte state embedded within a multicellular glial niche. The spatial localization of MCN2 and stGP4 further supported this correspondence, with both signals concentrated in deep white matter regions (Fig. 4a,d). Differential expression between aged and young oligodendrocytes provided an independent view of this decomposition. The strongest age-increased genes included shared MCN2 genes, such as *PDIA2, SORCS2, MAT2A* and *RGS11* (Fig. 4e), which were also recovered as high-weight genes in shared MCN2 and stGP4. Together with the increasing temporal effect of stGP4 (Fig. 4c), these results indicate that the molecular identity of this niche-related program is shared across sections, while its activity changes with aging. This provides empirical support for the stGP modeling assumption that gene programs can be shared, whereas their temporal effects and spatial deployment vary across biological conditions (Fig. 1).

In contrast to the stGP4-shared MCN2 niche, several strongly age-decreased genes were assigned to other shared MCNs, including SLC17A7, SLC1A2 and NEUROD6 (Fig. 4e). These genes were not part of the MCN2-stGP4 axis. Instead, they were more closely related to a distinct neuron-proximal program represented by stGP1, associated with NicheScope shared MCN1 genes (Supplementary Fig. S11). We further found that stGP1 activity was negatively correlated with distance to the nearest neuron, whereas stGP4 activity showed mostly positive correlations with nearest-neuron distance (Supplementary Fig. S11). Thus, the age-decreased MCN1-like genes define a neuron-proximal contrast to stGP4, rather than evidence for loss of the MCN2-associated oligodendrocyte niche.

NicheScope also detected young- and aged-specific MCNs, but these age-specific MCN genes contributed relatively few of the strongest age-associated signals in oligodendrocytes (Fig. 4e and Supplementary Fig. S12). Instead, the dominant age-associated changes were concentrated in shared MCN gene sets, with MCN2 genes increasing and other shared MCN genes decreasing. This pattern supports a model in which aging remodels the activity and anatomical deployment of pre-existing multicellular niches, rather than simply creating or eliminating age-specific niches. Together, these results show that stGP4 represents an age-varying, niche-coupled oligodendrocyte program, linking donor age to spatially localized remodeling of multicellular glial organization in the human DLPFC.

### stGP identifies a late-life, T-cell-associated microglial MHC-I program in aging white matter

To test whether stGP could generalize beyond human DLPFCs, we applied stGP to a mouse brain lifespan MERFISH dataset comprising 20 coronal sections^4^. We focused on microglia because brain aging is accompanied by spatially localized neuroimmune remodeling, particularly in white-matter regions^25-27^. stGP resolved four microglial gene programs with distinct contributions from age-related and within-section spatial variance components (Fig. 5b). Among them, stGP1 and stGP4 were dominated by within-section spatial variation (Supplementary Fig. S14), whereas stGP3 was almost entirely explained by the aging component (Supplementary Fig. S15). By contrast, stGP2 was the spatiotemporally variable program for microglia and was therefore selected in the remainder of the analysis. The aging effect of stGP2 increased progressively across the lifespan, with the steepest rise in late life (Fig. 5a). High-loading genes in stGP2 included core components of the major histocompatibility complex class I (MHC-I) molecule antigen-presentation machinery, such as *B2m, H2-K1*, and *H2-D1*^28^, together with interferon-responsive and immune-regulatory genes including *Bst2, Ifi27, Lag3* and *Pdcd1*^4,27,29^ (Supplementary Fig. S13). Over-representation analysis of active stGP2 genes further linked this program to a coherent MHC-I antigen-presentation axis, including antigen processing and presentation, peptide-loading machinery, T-cell-related binding or cytotoxicity terms, and a brain myeloid/microglial aging signature (Fig. 5f). Together, these results identify stGP2 as an aging-associated microglial immune program characterized by MHC-I antigen presentation and interferon-responsive expression, with increasingly elevated activity at older ages.

**Fig. 5.**
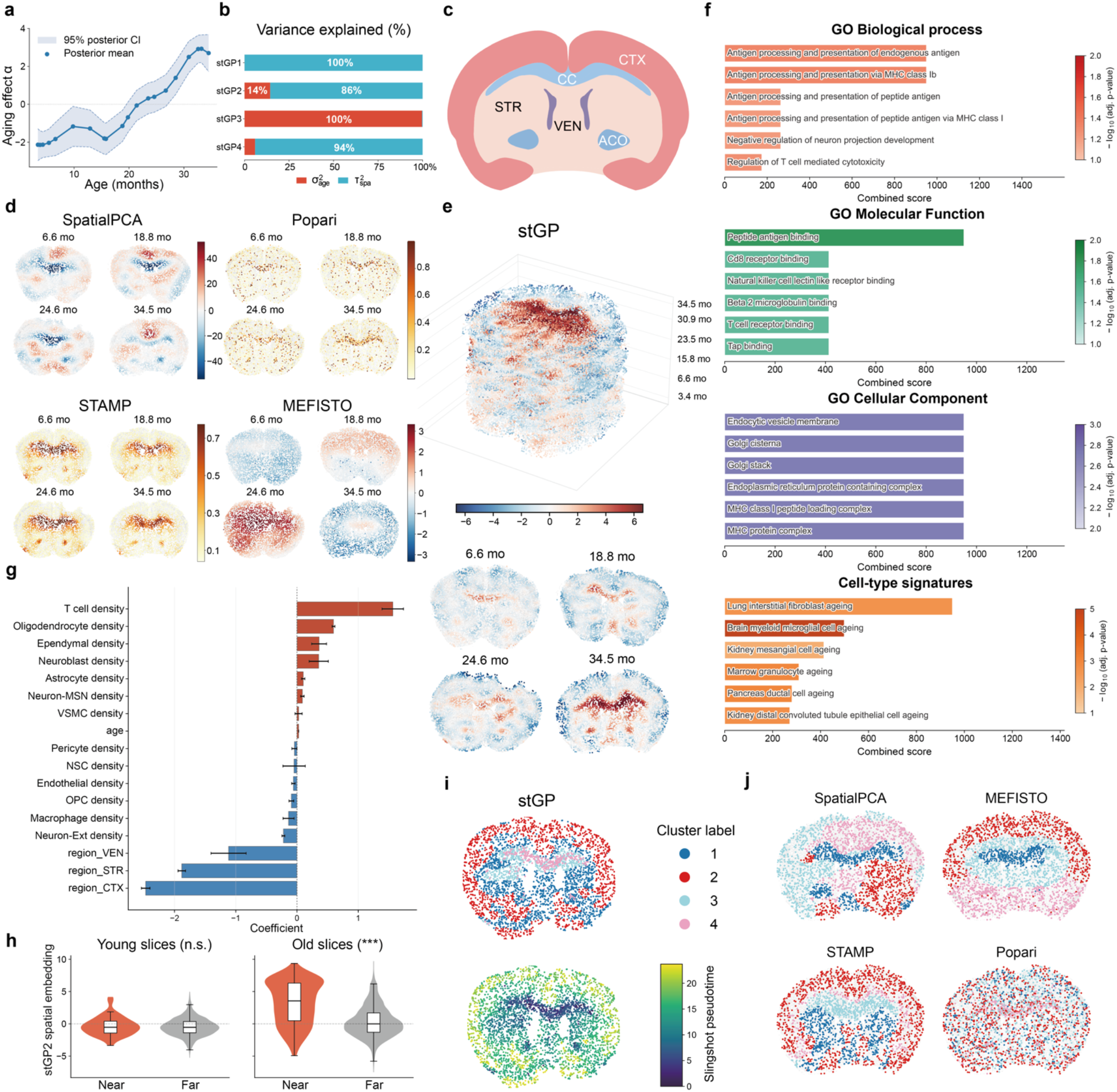
stGP identifies a late-life, T-cell-associated microglial MHC-I program in aging mouse white matter. **a**, The temporal effect trajectory of the stGP2 program in microglia. **b**, Spatial and temporal variance contributions across stGP programs. **c**, The anatomical regions in coronal mouse brain sections. **d**, The spatial embeddings produced by baseline matrix factorization methods. Top left: SpatialPCA; top right: Popari; bottom left: STAMP; bottom right: MEFISTO. **e**, The spatial activities of stGP2 across all 20 slices (top panel) and four representative slices (bottom panel). **f**, Functional enrichment of stGP2 genes, highlighting MHC-I antigen-presentation and interferon-responsive immune terms. **g**, The regression coefficients of stGP2 embeddings on local cell type densities, after controlling for age and region effects. **h**, The boxplots of stGP2 spatial activities on cells of near versus far from T cells, in both young and aged slices with two-sided Mann-Whitney U tests (*, *P* < 0.05; **, *P* < 0.01; ***, *P* < 0.001). **i**, The clustered spatial domains using stGP spatial embeddings (top panel) and the corresponding pseudotime inference by Slingshot (bottom panel). **j**, The clustered spatial domains using baseline methods. Top left: SpatialPCA; top right: MEFISTO; bottom left: STAMP; bottom right: Popari.

Spatially, stGP2 was not uniformly distributed across the brain. Instead, the spatial activity of stGP2 became enriched in white-matter tracts, with strongest localization to the corpus callosum (CC) and anterior commissure (ACO) at older ages (Fig. 5c,e and Supplementary Fig. S16). At young ages, stGP2 displayed only weak and patchy spatial structure, whereas aged sections showed a sharply focused CC/ACO-enriched pattern (Fig. 5e). Thus, stGP2 captures not only a global aging-associated increase (Fig. 5a), but also the dynamic spatial deployment of a shared microglial program across ages. This white-matter-enriched pattern is consistent with prior reports that white-matter tracts are hotspots of glial and immune remodeling during brain aging^25,26^, while extending those observations by resolving a microglia-specific program with interpretable gene loadings and separable temporal and spatial components.

Notably, this joint spatial–temporal interacted structure was not recovered by the baseline methods. SpatialPCA recovered spatially coherent factors but was confounded with temporal effect, which showed stable anatomical and patchy regions rather than the late-life CC/ACO-specific activation captured by stGP2 (Fig. 5d,e). Popari, although identified the CC/ACO region in very aged slices, still produced high and diffused microglial scores in cortex region, which did not isolate the white-matter related pattern in this cell-type-specific setting (Fig. 5d). MEFISTO modeled age and spatial coordinates through a single GP kernel and captured overly smooth variation without resolving the localized CC/ACO microglial program (Fig. 5d). STAMP identified a CC-enriched spatial topic, but failed to recover the broader white-matter program including ACO, which was together identified as the aging hotspot^27^, and the topic did not show the late-life increase observed for stGP2 (Fig. 5d, Supplementary Fig. S17). In terms of clustering the spatial domains, stGP produced coherent domains aligned with not only white matter but also regions like cortex (CTX) and striatum (STR) (Fig. 5c,i), whereas baseline methods were often dominated by broad anatomy, patchy patterns or spatially diffuse assignments (Fig. 5j). Together with the pseudotime inference results confirming the aging hotspot of white matter^25^, we found that stGP not only identified the stable tissue structure of mouse brains (Fig. 5i), but also revealed the dynamic enhancement of the spatially localized aging effect (Fig. 5e). These comparisons highlighted the importance of explicitly decomposing a shared temporal trajectory from section-specific spatial embeddings.

We next asked whether the white-matter activation of stGP2 reflected a local cellular niche beyond regional anatomy. Although stGP is fitted within microglia, it should also reflect the effects from local neighborhoods, since the spatial kernel of stGP operated at a local cellular neighborhood scale (Supplementary Fig. S19). Note that the spatial activity of stGP2 was highly colocalized with T cells, but cell type proportions also covary with regions (Supplementary Fig. S18). We therefore regressed the stGP2 embedding on local cell-type densities while adjusting for age and region indicators, finding that T-cell density emerged as the strongest positive predictor of stGP2 activities (Fig. 5g). Oligodendrocyte and ependymal densities also had positive coefficients, consistent with white-matter and ventricular-adjacent localization. Since temporal effects have been separated out by stGP2 temporal effect *α*, age did not have a significant effect on the stGP spatial embeddings. Conversely, cortical, striatal and ventricular region indicators were negative relative to the white-matter reference, indicating that anatomical region explained part but not all the spatial variations (Fig. 5g). This association is consistent with previous reports showing that T cells accumulate in aged neurogenic and white-matter environments and can promote interferon-responsive microglial states during brain aging^4,27,30^. A complementary matched near-versus-far analysis supported the age dependence of the T-cell association. Microglia located near T cells did not show higher stGP2 spatial activities than those far from T cells in young sections, whereas near-T-cell microglia in old sections showed a pronounced increase in stGP2 activities relative to those microglia farther from T cells (Fig. 5h). Thus, the late-life amplification of stGP2 is coupled to local T-cell-enriched neighborhoods after adjustment for regional anatomy, rather than reflecting white-matter identity alone as an aging hotspot.

### stGP reveals replicable and robust stage-specific repairing programs in the mouse injured kidney Xenium dataset

To demonstrate the effectiveness and reproducibility of stGP in a dynamic injury–repair setting, we analyzed a mouse kidney spatial transcriptomic time course generated after bilateral ischemia–reperfusion injury (BIRI)^7^. The original study profiled kidneys across sham, 4 hours, 12 hours, 2 days, 14 days, and 6 weeks after injury using high-resolution Xenium in situ measurements. Here, we focused on the five post-injury time points and fitted stGP independently to left and right kidney cohorts, treated as bilateral biological replicates. Including sham as the initial baseline produced qualitatively similar results (Supplementary Fig. S20). In immune cells, stGP identified three temporally ordered programs consistently across bilateral kidneys (Fig. 6d). Matching programs by cosine similarity of their gene loading showed a nearly one-to-one correspondence between the left and right kidney fits (Fig. 6b,e), suggesting the reproducibility of stGP. Sensitivity analyses also showed that the matched programs were stable across various kernel and bandwidth choices (Fig. 6g, Supplementary Fig. S21), supporting the robustness of the stGP method.

**Fig. 6.**
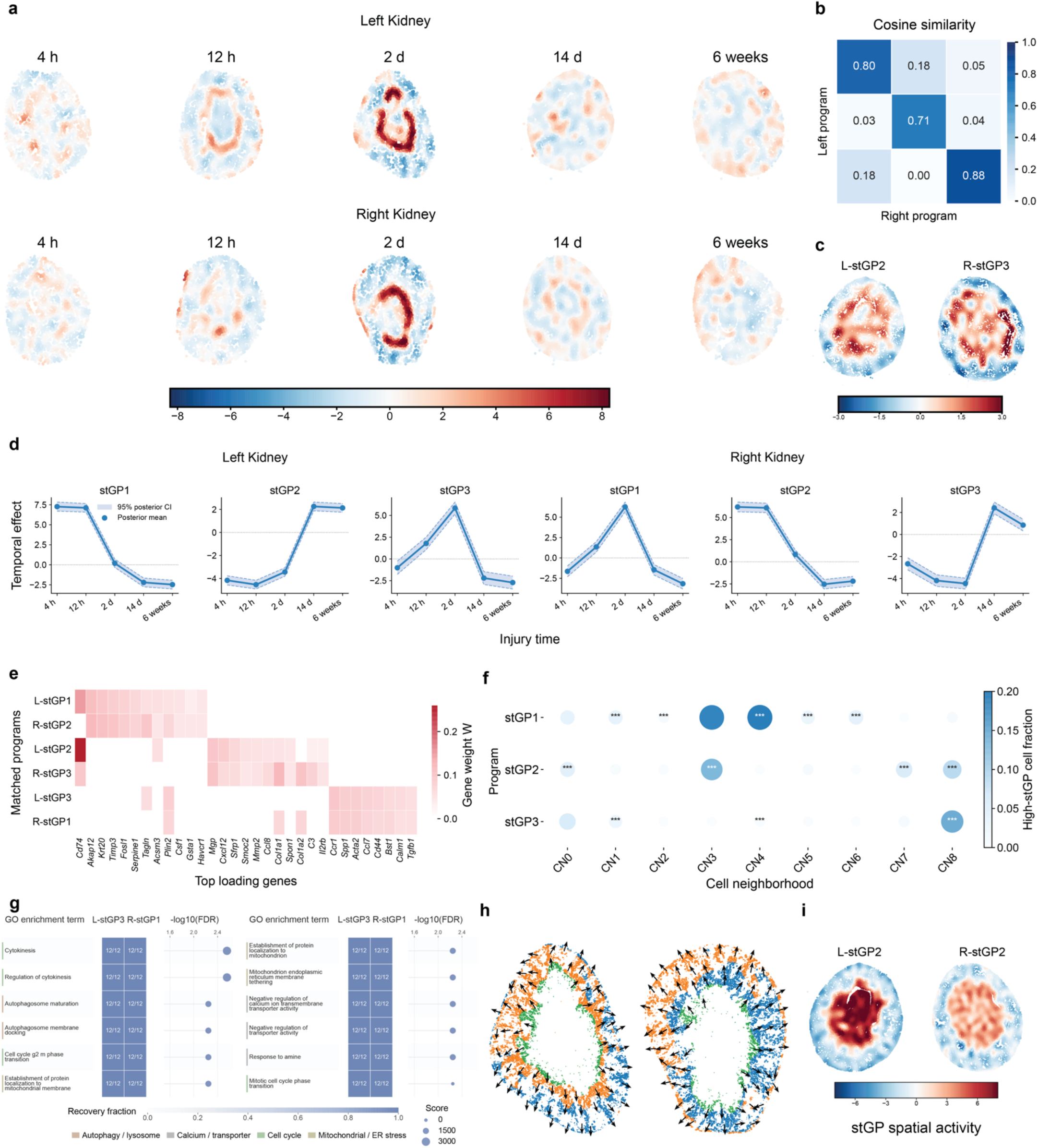
stGP reveals reproducible stage-specific repairing programs in the mouse BIRI kidney Xenium time course across bilateral kidneys. **a**, Spatial activities of the matched day-2 immune repair-transition programs learned from independent left-kidney (top panel, L-stGP3) and right-kidney (bottom panel, R-stGP1) immune cells. **b**, Cosine similarity between immune-program gene-loading vectors from independent left- and right-kidney stGP fits. **c**, Day-14 spatial activities of the matched late immune-stromal remodeling programs, L-stGP2 and R-stGP3. **d**, Temporal effects for the three immune programs in left and right kidney fits. **e**, Gene-loading heatmap for the matched left- and right-kidney immune stGP programs. **f**, Distribution of high-scoring immune cells across cellular neighborhoods (CNs) in the left-kidney immune fit. Dot color and size denote the fraction of cells in each CN that were high scoring for the corresponding program. Asterisks denote one-sided Fisher exact tests for enrichment of high-scoring cells in a CN versus all other CNs, adjusted by the Benjamini-Hochberg procedure (*, adj. *P* < 0.05; **, adj. *P* < 0.01; ***, adj. *P* < 0.001). **g**, Recurrent Gene Ontology (GO) Biological Process enrichments for the matched day-2 immune programs, L-stGP3 and R-stGP1, across 12 temporal/spatial kernel settings and bilateral cohorts. Heatmap entries indicate the number of settings in which each term was recovered. Dots show the enrichment significance in the representative setting, with point size indicating the Enrichr combined score. **h**, Pseudotime trajectories inferred from stGP spatial domains in day-2 injured proximal tubule cells across bilateral kidneys. Domains were obtained by spectral clustering of the stGP spatial embedding. Arrows indicate the inferred trajectory direction starting from the corticomedullary and outer-medullary injury zone. **i**, Day-14 spatial activities of fibroblast programs in left and right kidneys (L-stGP2 and R-stGP2).

The first immune program represented an acute injury-associated state. In the left kidney, L-stGP1 was already high at 4 hours and 12 hours after BIRI and declined by day 2 (Fig. 6d); the right-kidney counterpart showed a similar early trajectory (Fig. 6d). This indicates that the immune program was activated much earlier than its abundance peak at 14 days (Supplementary Fig. S22), suggesting that stGP separates molecular activation within immune cells from later expansion of the immune compartment. The second L-stGP3/R-stGP1 program peaked at day 2 and captured an early inflammatory-to-repair transition (Fig. 6d). This paired program showed high loadings for *Ccr1, Spp1, Ccl7, Cd44, Bst1* and *Tgfb1* (Fig. 6e), consistent with an activated inflammatory-repair state involving chemokine-mediated leukocyte recruitment, cell adhesion and tissue-remodeling responses after injury^31-33^. Coherent enrichment analysis further supports this program associated with immune-cell turnover and intracellular stress remodeling during early repair (Fig. 6g). Importantly, this program was not uniformly activated across the tissue. Beyond the slice-level temporal effect activated at day 2 (Fig. 6d), spatially, L-stGP3 and R-stGP1 exhibited a restricted annular band in both kidneys (Fig. 6a), anatomically aligned with the corticomedullary and outer-medullary injury zone (Fig. 6h)^7^. This indicates that stGP captured a spatiotemporally organized inflammatory repair response rather than a purely global temporal effect (Supplementary Fig. S23). The third immune program marked late immune-stromal remodeling during maladaptive repair^34^. L-stGP2 and R-stGP3 were sharply activated at day 14 and remained positive at 6 weeks (Fig. 6d). Their spatial activities were patchy and localized to cortical remodeling domains at day 14 (Fig. 6c), but largely resembling the independently learned fibroblast stGP activity at the same stage (Fig. 6i), suggesting coordinated immune–fibroblast remodeling rather than independent immune activation alone. Together, these three immune programs reconstruct a temporal transition from acute epithelial injury sensing, through inflammatory immune activation, to persistent immune–stromal coupling in the fibrotic kidney.

We next anchored the stGP immune programs to the existing cellular neighborhoods (CNs) defined by clustering cells according to local neighboring cell-type proportions^7^. Consistent with the spatial overlap between late immune and fibroblast stGP activities at day 14 (Fig. 6c,i), the late immune–stromal remodeling programs were concentrated in tubulointerstitial remodeling domains involving medullary proximal-tubule, fibro-inflammatory and uro-immune neighborhoods, corresponding to CN3, CN7 and CN8 (Fig. 6f, Supplementary Fig. S24). Among these, CN7 specifically denotes a fibro-inflammatory niche, providing an independently annotated spatial context for maladaptive repair and chronic fibrotic remodeling (Supplementary Fig. S24). In contrast, the acute program L-stGP1/R-stGP2 was enriched in early niches CN3 and CN4 (Fig. 6f), corresponding to medullary proximal tubule and injured proximal-tubule neighborhoods, respectively (Supplementary Fig. S24). This localization supports the acute-injury interpretation of L-stGP1 and links it to vulnerable medullary/S3 proximal-tubule regions after BIRI.

Finally, we used stGP to resolve spatially organized epithelial-state heterogeneity among day-2 injured proximal tubule (PT) cells, since day-2 injured proximal tubule cells were reported to follow divergent trajectories, with some cells returning toward healthy states and others transitioning toward failed-repair (FR) proximal tubule cells by day 14^7^. After fitting stGP, day-2 injured proximal tubule cells separated into three spatially organized domains in both side kidneys (Fig. 6h). Comparing these domains with the original injured PT state annotations (Inj_S1, Inj_S2, and Inj_S3)^7^, we found that a recovery/metabolic domain was dominated by Inj_S2 cells with higher marker scores, whereas the maladaptive pre-failed-repair domain near the corticomedullary and outer-medullary injury zone showed elevated acute-injury and maladaptive/pre-FR marker scores, enrichment for Inj_S1/Inj_S3 cells and the strongest proximity to fibroblast–immune neighborhoods (Supplementary Fig. S25). The remaining domain showed intermediate marker and neighborhood profiles, consistent with an acute-stress transitional state. Thus, the stGP-derived epithelial embedding demonstrated the divergent injured-proximal-tubule cell fates, separating cells with metabolic recovery features from cells entering a fibro-inflammatory, maladaptive pre-failed-repair niche.

Together, the immune, fibroblast and epithelial analyses show that stGP decomposes the kidney injury-repair process into reproducible and robust spatiotemporal programs spanning acute epithelial stress, inflammatory activation and persistent immune–stromal remodeling.

## Discussion

Spatiotemporal transcriptomic datasets increasingly make it possible to observe how molecular programs vary across biological time within spatially organized tissues. However, interpreting these data remains challenging because sample-level temporal progression, section-specific anatomy, and local niche-associated variation are observed together and can confound one another. Here, we developed stGP, a statistical framework for identifying interpretable cell-type-specific gene programs in multi-sample spatiotemporal transcriptomic studies. By decomposing per-cell program activity into a sample-level temporal trajectory and a within-section spatial effect while preserving shared non-negative gene loadings, stGP resolves whether a cell-type-specific program represents a shared temporal response, a stable anatomical structure, or the localized deployment of a temporal process within specific tissue microenvironments. This allows stGP to distinguish preserved tissue architecture from age-associated remodeling and to reveal when broad temporal programs become spatially restricted to anatomical domains, aging hotspots, or multicellular niches. Comprehensive simulations show that stGP accurately recovers gene programs and separates temporal from spatial variation with favorable computational efficiency. Applications to human DLPFC, aging mouse brain and injured mouse kidney further demonstrated its ability to resolve spatial domains, uncover niche-associated aging programs, and reconstruct replicable repair trajectories across biological replicates. These results support stGP as a versatile and interpretable framework for studying biological processes in which molecular state, spatial organization and biological time are coupled together.

Finally, we discuss here the limitations and potential extensions of stGP. First, although stGP could be efficient for various single-cell resolution spatial technologies (e.g., MERFISH and Xenium), it remains challenging to directly apply stGP to low-resolution, spot-based ST platforms such as 10x Visium for cell-type-specific analysis. Integrating single-cell references in a unified probabilistic framework^35^ should be useful for the versatile modeling of ST data across different technologies. Second, stGP is currently fitted within a prespecified target cell type. Although downstream analyses associate stGP programs with neighboring cell types and multicellular niches, the expression profiles or densities of other cell types are not explicitly included as covariates in the model. Future extensions could incorporate local cell-type composition or niche covariates into the model for program activities, thereby separating intrinsic spatiotemporal dynamics from the microenvironmental effects^36,37^. Third, stGP currently analyzes independent two-dimensional tissue sections. However, cells are organized within three-dimensional tissue architectures, and preserving this 3D spatial context can be important for deciphering complex biological processes. Extending stGP to reconstructed or directly profiled 3D tissues across biological time would broaden its applicability to volumetric spatial transcriptomic atlases and enable the analysis of spatiotemporal programs^38-40^.

In summary, stGP provides a versatile statistical framework for characterizing dynamic tissue architecture through cell-type-specific spatiotemporal gene programs. Across simulations and applications to human brain aging, mouse brain aging, and kidney injury repair, stGP resolves interpretable molecular programs, deciphers temporal and spatial sources of variation, and links program activity to local tissue organization. These features make stGP a useful tool for studying biological processes in which molecular state, spatial context, and biological time are tightly intertwined.

## Methods

### The stGP method

stGP is intended for multi-sample spatial transcriptomic data in which a target cell type is observed across samples with sample-level temporal covariates. The input for each cell type consists of an expression matrix, sample labels, temporal covariates, and within-section spatial coordinates. The outputs are gene program loadings, sample-level temporal effects, within-section spatial activities, and variance component estimates. The model was fitted independently within each cell type.

Let *t* = 1, …, *T* index biological samples or individuals at different time points. For each sample *t*, let 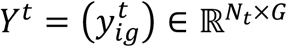 denote the *N*_*t*_ × *G* expression matrix for *N*_*t*_ cells and *G* genes in the target cell type. Let *μ* ∈ ℝ^*G*^ denote the baseline gene effect for this cell type. Assume that the *G* genes are represented by *p* gene programs. This yields a factor matrix *W* = (*w*_*jg*_) ∈ℝ_+_^p×*G*^ whose rows lie on the simplex^14^,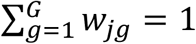.

This constraint fixes the scale of each loading vector and makes the gene programs directly interpretable as sparse or approximately sparse weighted gene sets. We then decompose the gene expression *Y*^*t*^ via the program matrix *W* and a spatiotemporal cell embedding matrix 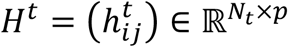. This leads to the following matrix decomposition:

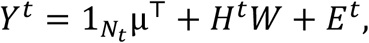

where 1_*N*_ = (1, …, 1)^⊤^ ∈ *R*^*N*^ denotes the all-one vector, and the noise term 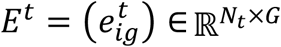 collects all the unexplained variation, with each entry 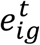 independently and identically follows a normal distribution with mean 0 and variance 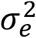. Without loss of generality, we remove the baseline effect in advance and hence write *Y*^*t*^ = *H*^*t*^ *W* + *E*^*t*^ in what follows. Specifically, we pre-center each gene across all cells of this cell type and therefore omit the intercept term *μ* as the intrinsic effect.

Note that there are many ways to define a cell embedding 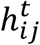. Here, we aim to learn a spatiotemporal embedding such that cells from samples with similar ages and nearby spatial locations within the same slice have similar embeddings. Specifically, for each program *j*, we decompose the per-cell activity 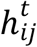 into a temporal effect α_*tj*_ and a within-section spatial component 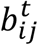 as:

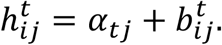

The sample-level term α_*tj*_ represents the temporal effect for program *j* in individual *t* and is shared by all cells of this target cell type within the sample. This is motivated by the synchronized temporal effects shared across cells from the individuals with the same chronological age^20,41^. Write α_*j*_ = (α_1*j*_, …, α_*Tj*_)^⊤^ ∈ ℝ^*T*^. We then assume that α follows a Gaussian process with mean 0 and covariance 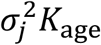, where 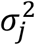 is the program-specific variance component. Here the temporal kernel *K*_age_ = (*K*_age,*tt’*_) ∈ ℝ^*T*×*T*^ captures the similarity between individuals with nearby ages, where

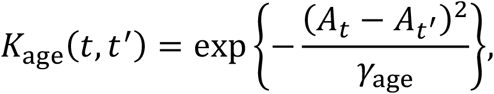

and *γ*_age_ is the temporal bandwidth that can be specified by the median technique (Supplementary Methods A.6). When the observed ages are sparse, we could apply the autoregressive kernel matrix 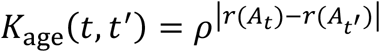 as an alternative, where *r*(⋅) is the ranking function. After considering the synchronizing temporal effect, 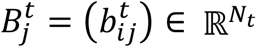 describes the heterogeneous spatial effect within section *t* on program *j*, since different programs may exhibit distinct expression patterns. We then assume 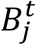 follows another Gaussian process with mean 0 and covariance 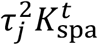, where 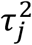 is the variance component for spatial effects^13^. The spatial kernel 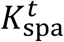 is defined separately within each section and does not require cross-section alignment:

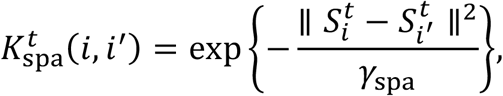

where 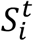 is the 2-dimensional spatial coordinate of the *i*-th cell of the *t*-th sample after standardization, and *γ*_spa_ is the spatial bandwidth. Because 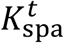 only depends on pairwise Euclidean distances within a section, the spatial prior is invariant to translation and rotation of each section^42^. The temporal and spatial effects, residual noise and program-specific priors were assumed independent across programs and samples.

### Numerical algorithm of stGP

We first describe the numerical algorithm of parameter estimation for a single program with *p* = 1. Stack cells across all samples so that *Y* is an *N* × *G* matrix, with *N* = ∑_*t*_ *N*_*t*_. For the rank-1 model, *Y* = *hw*^⊤^ + *E*. Let 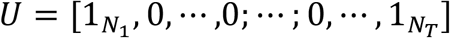 be the *N* × *T* block design matrix that maps each cell to its sample, such that (*U*α)_*i*_ = α_*t*_ for cells in sample *t*. Write *b* = ((*b*^1^)^⊤^, …, (*b*^*T*^)^⊤^)^⊤^ ∈ ℝ^*N*^, the latent embedding is *h* = *U*α + *b* with covariance

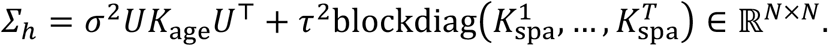

Note that the unknown parameters to be estimated are 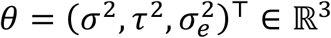 and the simplex-constrained loading vector *w* ∈ ℝ^*G*^, as well as the latent variable *h* ∈ ℝ^*N*^. stGP uses a blockwise procedure that alternates among a posterior update for *h*, a simplex-constrained update for *w* and a variance-component MM update for *θ* as follows.

First, when *w* and *θ* are given, the conditional posterior of *h* follows a Gaussian distribution with covariance 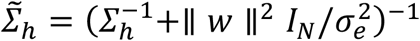 and mean 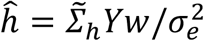, where 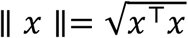 is the usual *ℓ*_2_-norm of an arbitrary vector *x*. Then, we use the MAP estimator 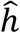 to replace *h*. Second, given the estimates of *h*, we update the gene program weights *w* under the simplex constraint 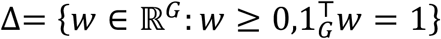 as 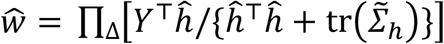. When exact sparsity is desired, the largest *K* coordinates of *w* can be retained before projection. Third, given the gene program *w*, we then update the variance components *θ* by maximizing the marginal log-likelihood of *y* = vec(*Y*). The rank-1 iteration continues until convergence (Supplementary Methods A.1-4).

For multiple programs with *p* > 1, stGP writes *Y* = *HW* + *E*, with *H* = [*h*_1_, …, *h*_*p*_] and with each row of *W* lying on the simplex. Cycling over *j* = 1, *⋯, p* with residuals 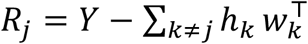, we apply the rank-1 updates to 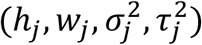 with *Y* replaced by *R*_*j*_. The model is then fitted by the backfitting algorithm^43^ to iteratively refine the estimates for each program given the estimates for the other programs. When the number of programs *p* is unknown, stGP first sets an upper bound *p*_E4F_ and automatically determines the number of components based on the component magnitude^44^ (Supplementary Methods A.6).

### Computational efficiency analysis

We analyze the computational complexity of the stGP method to assess its scalability. Recall that *T* denotes the number of samples, *N*_*t*_ is the number of cells in sample *t*, 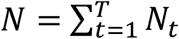 records the total number of cells, *G* refers to the number of genes. We focus on a single program at a time since the *p* programs can be updated by the blockwise backfitting algorithm. A direct implementation would require repeated inversion of the *N* × *N* latent covariance *Σ*_*h*_ for each program and inversion of an even larger *NG* × *NG* marginal covariance in the variance component MM update. These operations would scale as *O* (*N*^3^) and *O*((*NG*)^3^), respectively. stGP avoids these bottlenecks by exploiting the block-diagonal structure of the within-section spatial kernel and the low-rank temporal term at the sample level, using the Woodbury identity^45^ and an orthogonal rotation in gene space. This leads to a much reduced 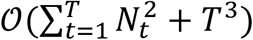 computation per update for both situations (Supplementary Methods A.2 and A.4).

We also recorded the computational time for stGP and the baseline methods in both simulation studies and real data analyses (Fig. 2e and Supplementary Table S1). The computation was performed on our Linux computing server equipped with an Intel Xeon Gold 6152 CPU @ 2.10 GHz.

## Downstream analysis

### Gene-program annotation and enrichment analysis

For each stGP fit, genes were ranked by their loading weights within each program. Over-representation analysis was performed with gseapy^46^ using the measured gene panel as the background. We used the Gene Ontology gene sets from the Molecular Signatures Database (MSigDB^47^) as our input gene sets. Enrichment p-values were adjusted for multiple testing within each analysis. In mouse kidney immune analyses, stGP programs were additionally tested against curated cell-neighborhood (CN) signatures^7^. For each program-CN pair, high-scoring cells were defined as the top 5% by stGP program score. CN enrichment was then tested using one-sided Fisher exact tests for enrichment of high-scoring cells within the CN and one-sided Mann-Whitney U tests comparing program scores inside versus outside the CN, followed by Benjamini-Hochberg adjustment.

### Proximity effect analysis

To test whether stGP program activity was associated with local cellular neighborhoods in aging mouse brains, we analyzed the within-section spatial activity *b* for each target cell after removing the shared temporal component as confounders. For each candidate neighboring cell type, target cells were classified as near if their nearest candidate cell was within 30 *μm* and far if the nearest candidate cell was more than 150 *μm* away within the same section. Near-versus-far effects were calculated as the difference in median *b* between near and far target cells. Adjusting for region effects as covariates, we regressed the stGP spatial embedding *b* on age, region indicators and log-transformed local cell-type densities within 50 *μm*. For an age-stratified validation, near-versus-far distributions of *b* were compared separately in young and aged sections using two-sided Mann-Whitney U tests.

### Niche analysis

For the human DLPFC analysis, multicellular niches associated with target cell types were identified using NicheScope^24^. Niche scores and niche-regulated cell-state genes were compared with stGP spatial embeddings and gene loadings to determine whether a program corresponded to a local multicellular microenvironment.

### Spatial domain clustering

To evaluate whether stGP spatial embeddings captured coherent spatial domains or cell niches, we performed graph-based clustering on the estimated within-section spatial embeddings. Specifically, each cell was represented by its stGP spatial embedding *b* after removing the temporal confounding *α*s. We constructed a *k*-nearest-neighbor (KNN) graph in the stGP embedding space using Euclidean distance. In benchmarking analyses with available reference labels, the number of clusters was set to the number of reference classes; in de novo analyses, the number of clusters was chosen according to biological or anatomical prior knowledge. The resulting clusters were referred to as stGP domains and were visualized on the original tissue coordinates to assess their spatial continuity and anatomical localization. When reference annotations were available (e.g., refined cell subtypes or marker genes), clustering performance could be evaluated using the adjusted Rand index (ARI)^22^ and normalized mutual information (NMI)^23^. Pseudotime trajectories were inferred using the stGP spatial domains obtained from the clustering analysis. A starting cluster was specified according to biological or anatomical prior knowledge in each dataset. Slingshot^48^ was then fitted to infer the pseudotime values, which were mapped back to the spatial coordinates for visualization.

### Benchmarking analysis

stGP was evaluated in simulation benchmarks and in real datasets. Simulation benchmarks compared stGP with PCA, NMF, SpatialPCA^13^, MEFISTO^17^, STAMP^16^ and Popari^15^. PCA and NMF^12^ provide non-spatial matrix factorization baselines; SpatialPCA is a spatially aware dimension reduction method; MEFISTO models smooth factors over observed covariates; STAMP is a spatial topic model with temporal smoothing; and Popari is a multi-sample spatial metagene model. Real-data benchmarks used the same filtered cell-type-specific matrices and matched ranks where possible. For signed methods, components were compared after sign-invariant transformation when non-negative program weights were required. See detailed settings in Supplementary Notes B.1.

### Differential expression

Differential expression analyses were used to compare genes associated with high-scoring program regions or age groups, as indicated in each analysis. Genes were tested within the relevant cell type, and P values were adjusted for multiple comparisons.

## Data resources and preprocessing

### Human aging brain MERFISH

We analyzed the human DLPFC MERFISH dataset^5^. Cells with missing or invalid cell-type annotations were removed. We excluded one flagged donor-region sample with apparent batch effects (donor 5823, rep1) and the 0.4-year infant sample before concatenating cohorts. Tissue sections were defined by donor-region identifiers, and donor age in years was used as the temporal covariate. Spatial coordinates were defined using recorded cell centroids. We only included the cells with following major cell types: astrocytes, endothelial cells, excitatory neurons, inhibitory neurons, microglia, oligodendrocytes and oligodendrocyte precursor cells. Finer annotations, including excitatory-neuron subtype, were retained where available for downstream layer-based evaluation of spatial domains. Cells were removed if they had fewer than 20 counts or fewer than 10 detected genes, exceeded the 99.9th percentile for counts or detected genes, had segmentation bounding-box areas outside the 0.1st-99.9th percentiles, or had anisotropy above the 99.5th percentile. To reduce likely neuronal contamination, non-neuronal cells with neuronal-marker transcript fraction >0.15 were removed, and non-neuronal cells within *μ*m of the nearest neuron in the same section were also removed.

### Mouse aging brain MERFISH

We analyzed the mouse aging brain MERFISH dataset^4^. The analysis was restricted to prespecified major neuronal, glial, vascular and immune cell populations. A curated set of high-spillover or misallocation marker genes from the original study was removed before downstream analysis. A segmentation bounding-box area was computed from min/max spatial coordinates. Cells were removed if they had fewer than 40 total counts or fewer than 15 detected genes, if their total counts or detected genes exceeded the 99.9th percentile, or if their bounding-box areas were outside the 0.1st-99.9th percentiles

### Mouse kidney injury and repair Xenium dataset

We analyzed the mouse BIRI kidney Xenium time course with 4 hours, 12 hours, 2 days, 14 days, and 6 weeks after injury^7^. Analyses were performed separately for left and right kidneys. This dataset also contains a sham control performed at day 14, which was dropped for the main analysis. Sham-inclusive immune analyses were run as a robustness analysis. For the stGP fits, expression data were already scaled before fitting, spatial coordinates were standardized within section. Different time points after injury were treated as different stages. We used an AR(1) temporal kernel over ordered post-injury stages with *ρ* = 0.7 for the stGP fits. Different values of *ρ* for the temporal kernel and *γ*_?)4_ for the spatial kernel were tested.

## Data availability

The aging mouse brain MERFISH dataset^4^ can be accessed from Zenodo at [https://doi.org/10.5281/zenodo.13883177]. The aging human brain DLPFC MERFISH dataset^5^ can be obtained from [https://publications.wenglab.org/SomaMut/]. The mouse kidney injury and repair Xenium dataset^7^ is available from [https://doi.org/10.6084/m9.figshare.28761695.v1].

## Code availability

The stGP software and scripts for reproducing the analyses are available at: https://github.com/YangLabHKUST/stGP.

## Acknowledgements

This work was supported in part by the Innovation and Technology Commission (ITC-SKLNSD26SC01); Hong Kong Research Grants Council Grants, AoE/E-601/24-N, C6040-24G, 16307221, 16307322, 16302823, 16309424, and 16308925; The Hong Kong University of Science and Technology Startup Grants R9405 and Z0428 from the Big Data Institute. The computation tasks for this work were partially performed using the X-GPU cluster supported by the Research Grants Council Collaborative Research Fund Grant C6021-19EF and HKUST SuperPOD.

## Author contributions

B.Y. and C.Y. conceived the study. B.Y. developed, implemented, and validated the stGP method. Z.T. helped with analyzing the results. B.Y. and C.Y. wrote the manuscript. Z.T., X.W., and H.W. provided critical feedback during the study and helped revise the manuscript.

