## Supplementary Information for "Characterizing dynamic tissue architectures by identifying cell-type-specific spatiotemporal gene programs with stGP"

**D Supplementary Tables** **44**

### A Supplementary Methods

#### A.1 The Rank-1 Model Setup

Recall that the rank-1 model for each individual $t = 1, \cdots, T$ is given by $Y^t = h^t w^\top + E^t$, where $E^t_{ig} \overset{\text{i.i.d.}}{\sim} N(0, \sigma_e^2)$, $Y^t \in \mathbb{R}^{N_t \times G}$, $h^t \in \mathbb{R}^{N_t}$, and $w \in \mathbb{R}^G$ is constrained to the simplex $\Delta = \{w \in \mathbb{R}^G : w \geq 0,\ 1_G^\top w = 1\}$. Let $N = \sum_{t=1}^T N_t$. Stack $Y = [(Y^1)^\top, \cdots, (Y^T)^\top]^\top \in \mathbb{R}^{N \times G}$ and $h = [(h^1)^\top, \cdots, (h^T)^\top]^\top \in \mathbb{R}^N$. In particular, the spatiotemporal embedding decomposes as $h^t = \alpha_t 1_{N_t} + b^t$, where $\alpha = (\alpha_1, \cdots, \alpha_T)^\top \in \mathbb{R}^T$ captures the across-individual temporal variation and $b^t \in \mathbb{R}^{N_t}$ captures within-sample spatial variation. Next, define the block design matrix $U \in \mathbb{R}^{N \times T}$ such that $(U\alpha)_i = \alpha_t$ for cell $i$ in individual $t$:

$$U = \begin{bmatrix} 1_{N_1} & 0 & \cdots & 0 \\ 0 & 1_{N_2} & \cdots & 0 \\ \vdots & \vdots & \ddots & \vdots \\ 0 & 0 & \cdots & 1_{N_T} \end{bmatrix},$$

and stack $b = [(b^1)^\top, \cdots, (b^T)^\top]^\top \in \mathbb{R}^N$, so that $h = U\alpha + b$. We place independent Gaussian process priors on the random effects as $\alpha \sim N(0,\ \sigma^2 K_{\text{age}})$ and $b \sim N(0,\ \tau^2 K_{\text{spa}})$, where $K_{\text{spa}} = \text{blockdiag}(K^1_{\text{spa}}, \cdots, K^T_{\text{spa}})$ is block diagonal. The induced prior on $h$ is $h \sim N(0, \Sigma_h)$ with $\Sigma_h = \sigma^2 U K_{\text{age}} U^\top + \tau^2 K_{\text{spa}}$. In what follows, we write $\theta = (\sigma^2, \tau^2, \sigma_e^2)^\top$ to collect the variance component parameters.

#### A.2 The MAP Update for $h$

In the following, we derive the maximum a posteriori (MAP) update for $h$.

##### A.2.1 The reduced model

Conditional on $(h, w)$, the log-likelihood is given by

$$\begin{aligned}\log p(Y \mid h, w, \sigma_e^2) &= -\frac{1}{2\sigma_e^2}\|Y - hw^\top\|_F^2 + C, \\ &= -\frac{1}{2\sigma_e^2}\Big\{\|Y\|_F^2 - 2h^\top(Yw) + \|w\|^2 h^\top h\Big\} + C, \end{aligned} \tag{A.1}$$

where $C$ is some constant independent of $h$. Therefore, for fixed $w$, the likelihood depends on $Y$ through the $N$-vector $Yw$ only. Define the normalized statistic $z = Yw/\|w\|^2 \in \mathbb{R}^N$ and its noise variance $\sigma_\eta^2 = \sigma_e^2/\|w\|^2$. Up to constants, (A.1) is equivalent to

$$\log p(Y \mid h, w, \sigma_e^2) = -\frac{1}{2\sigma_\eta^2}\|z - h\|^2 + C',$$

which can be interpreted as the pseudo-observation reduced model $z = h + \eta$, where $\eta \sim N(0, \sigma_\eta^2 I_N)$. This is an exact linear reduction given $w$, which is used to simplify algebra and computation as follows.

##### A.2.2 Posterior of $h$ given $(w, \theta)$

Combining the Gaussian prior $h \sim N(0, \Sigma_h)$ with (A.1), we obtain the Gaussian posterior $h \mid (Y, w, \theta) \sim N(\widehat{h},\ \widetilde{\Sigma}_h)$ with

$$\begin{aligned}\widetilde{\Sigma}_h &= \Big(\Sigma_h^{-1} + \frac{\|w\|^2}{\sigma_e^2} I_N\Big)^{-1}, \\ \widehat{h} &= \widetilde{\Sigma}_h\Big(\frac{Yw}{\sigma_e^2}\Big) = \widetilde{\Sigma}_h\Big(\frac{z}{\sigma_\eta^2}\Big). \end{aligned} \tag{A.2}$$

This posterior mean $\widehat{h}$ is then used for the MAP update in the rank-1 update procedure.

Next, note that $\Sigma_h$ and $\widetilde{\Sigma}_h$ are $N \times N$ matrices, whose inverse takes $O(N^3)$ computation. To address this issue, we rewrite the posterior mean in (A.2) as the following linear system

$$(I_N + \kappa\Sigma_h)\widehat{h} = \xi,$$

where $\kappa = \|w\|^2/\sigma_e^2$ and $\xi = \Sigma_h\big(Yw/\sigma_e^2\big)$. Recall that

$$I_N + \kappa\Sigma_h = \big(I_N + \kappa\tau^2 K_{\text{spa}}\big) + U\big(\kappa\sigma^2 K_{\text{age}}\big)U^\top = A + UMU^\top,$$

where $A = \text{blockdiag}(A_1, \cdots, A_T)$ is a block-diagonal matrix with $A_t = I_{N_t} + \kappa\tau^2 K_{\text{spa}}^t$, and $UMU^\top = U\big(\kappa\sigma^2 K_{\text{age}}\big)U^\top$ is a low-rank matrix with rank at most $T$. Therefore, by the Woodbury identity (Woodbury, 1950), we have

$$\begin{aligned}(I_N + \kappa\Sigma_h)^{-1} &= (A + UMU^\top)^{-1} \\ &= A^{-1} - A^{-1}U\big(M^{-1} + U^\top A^{-1}U\big)^{-1}U^\top A^{-1}.\end{aligned}$$

Following standard eigendecomposition techniques, the posterior mean $\widehat{h}$ derived in (A.2) can be computed efficiently with $O\big(\sum_{t=1}^T N_t^2 + T^3\big)$ cost. Similarly, $\text{tr}\big(\widetilde{\Sigma}_h\big)$ can also be computed without forming any $N \times N$ dense matrix.

##### A.2.3 Posterior distribution of $\alpha$ and $b$ given $(w, \theta)$

Using the pseudo-observation reduced model, we obtain a linear Gaussian system

$$z = U\alpha + b + \eta, \qquad \eta \sim N(0, \sigma_\eta^2 I_N),$$

with independent priors on $\alpha$ and $b$. Let $D = \tau^2 K_{\text{spa}} + \sigma_\eta^2 I_N = \text{blockdiag}(D_1, \cdots, D_T)$ with $D_t = \tau^2 K_{\text{spa}}^t + \sigma_\eta^2 I_{N_t}$. Then $\Sigma_z = \text{Cov}(z \mid w, \theta) = \sigma^2 U K_{\text{age}} U^\top + D$.

Standard Gaussian conditioning gives the posterior mean of $\alpha$ and $b$ as

$$\mathbb{E}[\alpha \mid z, w, \theta] = \sigma^2 K_{\text{age}} U^\top \Sigma_z^{-1} z,$$

$$\mathbb{E}[b \mid z, w, \theta] = \tau^2 K_{\text{spa}} \Sigma_z^{-1} z.$$

Consequently, we have

$$\mathbb{E}[h \mid z, w, \theta] = U\,\mathbb{E}[\alpha \mid z, w, \theta] + \mathbb{E}[b \mid z, w, \theta] = \Sigma_h \Sigma_z^{-1} z,$$

which matches the posterior mean $\widehat{h}$ in (A.2) because $\Sigma_z = \Sigma_h + \sigma_\eta^2 I_N$.

Next, by the Bayes rule, we further obtain the joint posterior distribution of $\alpha$ and $b$ as

$$\begin{bmatrix} \alpha \\ b \end{bmatrix} \Bigg| \; z, w, \theta \sim N\left( Q^{-1} \begin{bmatrix} \sigma_\eta^{-2} U^\top z \\ \sigma_\eta^{-2} z \end{bmatrix}, Q^{-1} \right),$$

where

$$Q = \begin{bmatrix} \sigma^{-2} K_{\text{age}}^{-1} + \sigma_\eta^{-2} U^\top U & \sigma_\eta^{-2} U^\top \\ \sigma_\eta^{-2} U & \tau^{-2} K_{\text{spa}}^{-1} + \sigma_\eta^{-2} I_N \end{bmatrix}.$$

Write the posterior covariance in block form as $Q^{-1} = [V_{\alpha\alpha}, V_{\alpha b}; V_{b\alpha}, V_{bb}] \in \mathbb{R}^{(T+N)\times(T+N)}$ with $V_{\alpha\alpha} = \text{cov}(\alpha \mid z, w, \theta) \in \mathbb{R}^{T\times T}$, $V_{bb} = \text{cov}(b \mid z, w, \theta) \in \mathbb{R}^{N\times N}$, and $V_{\alpha b} = \text{cov}(\alpha, b \mid z, w, \theta) \in \mathbb{R}^{T\times N}$. We then have the marginal posterior covariance of $\alpha$ given by

$$\begin{aligned} V_{\alpha\alpha} = \text{cov}(\alpha \mid z, w, \theta) &= \sigma^2 K_{\text{age}} - \sigma^4 K_{\text{age}} U^\top \Sigma_z^{-1} U K_{\text{age}} \\ &= \left(\sigma^{-2} K_{\text{age}}^{-1} + U^\top D^{-1} U\right)^{-1}. \end{aligned}$$

For uncertainty quantification, we use the plug-in posterior distribution at the fitted values $(\widehat{w}, \widehat{\theta})$ to obtain $\widehat{V}_{\alpha\alpha}$. In particular, for an arbitrary confidence level $1 - q$, a pointwise posterior interval for the aging effect $\alpha_t$ is constructed as

$$\widehat{\alpha}_t \pm z_{1-q/2}\sqrt{\left(\widehat{V}_{\alpha\alpha}\right)_{tt}},$$

where $z_{1-q/2}$ is the $(1-q/2)$-th quantile of the standard normal distribution. In the main text, we report these pointwise posterior intervals with $q = 0.05$.

#### A.3 Uncertainty-aware Update for $w$

The update for $w$ is analogous to the $M$-step in the EM algorithm (Dempster et al., 1977), with the additional simplex constraint. Recall that in the previous section, we derived the posterior distribution of the latent variable $h$ as $h \mid (Y, w, \theta) \sim N(\widehat{h}, \widetilde{\Sigma}_h)$. We update $w$ by minimizing the posterior expected squared error:

$$\widehat{w} = \arg\min_{w \in \Delta} \mathbb{E}_h \left\{ \|Y - hw^\top\|_F^2 \;\middle|\; Y, w^{(k)}, \theta^{(k)} \right\},$$

where $\Delta = \{w \in \mathbb{R}^G : w \geq 0,\ 1_G^\top w = 1\}$ is the simplex constraint and $\|A\|_F$ refers to the Frobenius norm. By taking the derivative with respect to $w$, we then obtain the updating formula of the unconstrained weight vector $w_{\text{unc}} = \left(Y^\top \widehat{h}\right) / \left\{\|\widehat{h}\|^2 + \text{tr}(\widetilde{\Sigma}_h)\right\}$. Projecting onto the simplex gives the update used in the algorithm:

$$\widehat{w} = \Pi_\Delta \left( \frac{Y^\top \widehat{h}}{\|\widehat{h}\|^2 + \text{tr}(\widetilde{\Sigma}_h)} \right).$$

Moreover, if exact $K$-sparsity is desired, we first keep the top-$K$ entries of $w_{\text{unc}}$ and then apply $\Pi_\Delta(\cdot)$ to the truncated vector, where $\Pi_\Delta(\cdot)$ denotes the Euclidean projection onto the simplex.

#### A.4 MM Updates for Variance Components $\theta$

Recall that $Y = hw^\top + E \in \mathbb{R}^{N\times G}$ collects all the gene expressions. With $w$ fixed, the marginal covariance of $y = \mathrm{vec}(Y) \in \mathbb{R}^{NG}$ is given by $\Omega(\theta) = \mathrm{cov}(y) = (ww^\top) \otimes \Sigma_h + \sigma_e^2 I_{NG}$. The marginal log-likelihood of $y$ is then given by

$$\mathcal{L}(\theta|y;w) = -\log|\Omega(\theta)|/2 - y^\top\Omega(\theta)^{-1}y/2.$$

By the convex property (Zhou et al., 2019), we can construct a surrogate function

$$g(\theta|\theta^{(k)}) = -\log\left|\Omega^{(k)}\right|\Big/2 - \mathrm{tr}\Big\{\Omega^{-(k)}(\Omega - \Omega^{(k)})\Big\}\Big/2$$
$$-y^\top\Omega^{-(k)}\Big\{\frac{\sigma^{4(k)}}{\sigma^2}ww^\top \otimes UK_{\mathrm{age}}U^\top + \frac{\tau^{4(k)}}{\tau^2}ww^\top \otimes K_{\mathrm{spa}} + \frac{\sigma_e^{4(k)}}{\sigma_e^2}I_{NG}\Big\}\Omega^{-(k)}y\Big/2,$$

where $\Omega^{(k)}$ refers to the covariance matrix $\Omega(\theta)$ with the $k$-step variance component estimator $\theta^{(k)} = (\sigma^{2(k)}, \tau^{2(k)}, \sigma_e^{2(k)})^\top$. It can be verified that $\mathcal{L}(\theta|y;w) \geq g(\theta|\theta^{(k)})$ for any $\theta$, with equality at $\theta = \theta^{(k)}$. Then, it suffices to iteratively maximize the surrogate function $g(\theta|\theta^{(k)})$ to maximize the marginal log-likelihood $\mathcal{L}(\theta)$.

Maximizing this surrogate function $g(\theta|\theta^{(k)})$ by taking the derivative with respect to $\theta$, we obtain the following updating formula from the first-order condition:

$$\sigma^{2(k+1)} = \sigma^{2(k)}\sqrt{\frac{y^\top\Omega^{-(k)}(ww^\top \otimes UK_{\mathrm{age}}U^\top)\Omega^{-(k)}y}{\mathrm{tr}\big\{\Omega^{-(k)}(ww^\top \otimes UK_{\mathrm{age}}U^\top)\big\}}}, \tag{A.3}$$

$$\tau^{2(k+1)} = \tau^{2(k)}\sqrt{\frac{y^\top\Omega^{-(k)}\{ww^\top \otimes K_{\mathrm{spa}}\}\Omega^{-(k)}y}{\mathrm{tr}\big[\Omega^{-(k)}\{ww^\top \otimes K_{\mathrm{spa}}\}\big]}}, \tag{A.4}$$

$$\sigma_e^{2(k+1)} = \sigma_e^{2(k)}\sqrt{\frac{y^\top\Omega^{-(k)}\Omega^{-(k)}y}{\mathrm{tr}\big(\Omega^{-(k)}\big)}}. \tag{A.5}$$

However, note that $\Omega$ is an $NG \times NG$ matrix, whose inverse is computationally infeasible. To address this issue, we apply the orthogonal matrix technique as follows. Specifically, let $\widetilde{w} = w/\|w\|$ and extend $\widetilde{w}$ to an orthonormal basis of $\mathbb{R}^G$, i.e., define $Q = [\widetilde{w},\ q_2,\ \cdots,\ q_G] \in \mathbb{R}^{G\times G}$ such that $Q^\top Q = I_G$ and $q_2, \cdots, q_G \in \mathbb{R}^G$ are orthogonal to $\widetilde{w}$. Next, define the orthogonally transformed vector $\widetilde{y} = (Q^\top \otimes I_N)y = \big(\widetilde{y}_1^\top, \cdots, \widetilde{y}_G^\top\big)^\top \in \mathbb{R}^{NG}$, where $\widetilde{y}_g \in \mathbb{R}^N$ and particularly $\widetilde{y}_1 = Y\widetilde{w} = Yw/\|w\| \in \mathbb{R}^N$. Moreover, since $w = \|w\|\widetilde{w} = \|w\|Qe_1$ with $e_1 = (1, 0, \cdots, 0)^\top \in \mathbb{R}^G$, we have $Q^\top ww^\top Q = \|w\|^2 e_1 e_1^\top$. Therefore, we then have the covariance matrix $\text{cov}(\widetilde{y}) = \widetilde{\Omega}$ as

$$
\begin{aligned}
\widetilde{\Omega} &= (Q^\top \otimes I_N)\Omega(Q \otimes I_N) \\
&= (Q^\top ww^\top Q) \otimes \Sigma_h + \sigma_e^2 I_{NG} \\
&= \text{diag}\Big(\underbrace{\|w\|^2\Sigma_h + \sigma_e^2 I_N}_{\Sigma_1},\ \sigma_e^2 I_N,\ \cdots,\ \sigma_e^2 I_N\Big).
\end{aligned}
$$

Then, in the MM updates, for the age variance component (A.3), we have the numerator as $y^\top\Omega^{-1}(ww^\top \otimes UK_{\text{age}}U^\top)\Omega^{-1}y = \|w\|^2\widetilde{y}_1^\top\Sigma_1^{-1}UK_{\text{age}}U^\top\Sigma_1^{-1}\widetilde{y}_1$, and the denominator $\text{tr}\big\{\Omega^{-1}(ww^\top \otimes UK_{\text{age}}U^\top)\big\} = \|w\|^2\text{tr}\big(\Sigma_1^{-1}UK_{\text{age}}U^\top\big)$. Since the structures of (A.4) and (A.5) are similar to (A.3), their computation can be treated very similarly and further simplified using the Woodbury identity and eigendecomposition as in Section A.2.2. This reduces the computational cost from $O((NG)^3)$ to $O(\sum_{t=1}^T N_t^2 + T^3)$.

#### A.5 Model Identifiability

In this subsection, we study the identifiability of the stGP model. Recall that, after centering the baseline gene effect, the multi-program stGP model can be written as

$$
Y = \sum_{j=1}^{p} h_j w_j^\top + E,
$$

where $Y \in \mathbb{R}^{N\times G}$ is the observed data matrix, $w_j \in \mathbb{R}_+^G$ is the $j$-th weight vector, and each row of $W = (w_1^\top, \cdots, w_p^\top)^\top \in \mathbb{R}_+^{p\times G}$ lies on the simplex $\Delta = \{w \geq 0 :\ 1_G^\top w = 1\}$. For the $j$-th program, the latent embedding $h_j \in \mathbb{R}^N$ is modeled as $h_j = U\alpha_j + b_j$, where $\alpha_j \sim N(0, \sigma_j^2 K_{\text{age}})$ and $b_j \sim N(0, \tau_j^2 K_{\text{spa}})$. Define $K_A = U K_{\text{age}} U^\top \in \mathbb{R}^{N\times N}$ and $K_S = K_{\text{spa}} \in \mathbb{R}^{N\times N}$. We then have $h_j \sim N\big(0, \Sigma_j\big)$, where $\Sigma_j = \sigma_j^2 K_A + \tau_j^2 K_S$. Write $C = \text{diag}(\sigma_1^2, \cdots, \sigma_p^2)$ and $D = \text{diag}(\tau_1^2, \cdots, \tau_p^2)$. Then we have $y = \text{vec}(Y) \sim N\big(0, \Omega\big)$, where $\Omega = \sum_{j=1}^p (w_j w_j^\top) \otimes \Sigma_j + \sigma_e^2 I_{NG}$. Equivalently, we have $\Omega = M_A \otimes K_A + M_S \otimes K_S + \sigma_e^2 I_{NG}$, where $M_A = W^\top C W = \sum_{j=1}^p \sigma_j^2 w_j w_j^\top$ and $M_S = W^\top D W = \sum_{j=1}^p \tau_j^2 w_j w_j^\top$. Note that the distribution of $y$ is only attributed to the covariance matrix $\Omega$. Therefore, the identifiability problem reduces to whether the map $(W, C, D, \sigma_e^2) \longmapsto \Omega$ is injective, regardless of the permutation of the $p$ programs.

It is worth emphasizing that the stGP model is not globally identifiable in complete generality. To see this, consider the case $p = 2$ with $\sigma_j^2, \tau_j^2 > 0$, and suppose the two programs share the same temporal-to-spatial variance ratio $\lambda_1 = \lambda_2 = \lambda$, where $\lambda_j = \sigma_j^2/\tau_j^2$ for $j = 1, 2$. Then, assume $L = D^{1/2} W \in \mathbb{R}_+^{2\times G}$ is strictly positive and has rank 2. Since $C = \lambda D$, we have $M_S = W^\top D W = L^\top L$ and $M_A = W^\top C W = \lambda W^\top D W = \lambda L^\top L$. Now let $R \in \mathbb{R}^{2\times 2}$ be an orthogonal matrix such that $\widetilde{L} = RL$ remains strictly positive. Next, write $\widetilde{L} = [\widetilde{l}_1, \widetilde{l}_2]^\top \in \mathbb{R}_+^{2\times G}$, where $\widetilde{l}_j^\top$ denote the $j$-th row of $\widetilde{L}$. Then, let $\widetilde{\tau}_j = \widetilde{l}_j^\top 1_G$, $\widetilde{\sigma}_j^2 = \lambda \widetilde{\tau}_j^2$, and $\widetilde{w}_j = \widetilde{l}_j / \widetilde{\tau}_j \in \mathbb{R}_+^G$. It can be verified that $\widetilde{w}_j$ still lies on the simplex. Moreover, with $\widetilde{D} = \text{diag}(\widetilde{\tau}_1^2, \widetilde{\tau}_2^2)$ and $\widetilde{C} = \text{diag}(\widetilde{\sigma}_1^2, \widetilde{\sigma}_2^2) = \lambda \widetilde{D}$, we obtain that

$$
\begin{aligned}
\widetilde{W}^\top \widetilde{D} \widetilde{W} &= \widetilde{L}^\top \widetilde{L} = L^\top R^\top R L = L^\top L = W^\top D W, \\
\widetilde{W}^\top \widetilde{C} \widetilde{W} &= \lambda \widetilde{L}^\top \widetilde{L} = \lambda L^\top L = W^\top C W.
\end{aligned}
$$

Hence $\widetilde{M_S} = M_S$ and $\widetilde{M_A} = M_A$, which means that the stGP model is not globally identifiable.

To fix this issue, we impose the following mild conditions to ensure identifiability, each of which has a natural biological interpretation.

**(C1)** *(Linear Independence of Covariance Components)* Assume that $K_A$, $K_S$, and $I_N$

are linearly independent.

**(C2)** *(Nonredundancy of Gene Programs)* Assume that $W$ has full row rank $p$, and that $\sigma_j^2 + \tau_j^2 > 0$ for all $j = 1, \cdots, p$.

**(C3)** Assume that at least one of the following two conditions holds.

**(C3a)** *(Variance Heterogeneity)* The variance proportions $\gamma_j = \sigma_j^2 / (\sigma_j^2 + \tau_j^2)$ are distinct for $j = 1, \cdots, p$.

**(C3b)** *(Program Separability)* For each program $j$, there exists a gene $g_j \in \{1, \cdots, G\}$ such that $w_{jg_j} > 0$ and $w_{kg_j} = 0$ for all $k \neq j$.

The above conditions are mild in the applications of interest. Condition (C1) requires that temporal smoothness, spatial smoothness, and white noise leave distinguishable signatures in the covariance structure (Hunter et al., 2021). This is easily satisfied, since the temporal covariance $K_A = UK_{\text{age}}U^\top$ has up to rank $T$, the spatial covariance $K_S$ is block-diagonal, and the white noise $I_N$ is fully diagonal. Condition (C2) rules out redundant latent programs and excludes the degenerate case in which a program contributes neither temporal nor spatial variation (Fu et al., 2018). Condition (C3a) requires different programs to have different temporal-to-total variance proportions. Condition (C3b) requires each program to have at least one sufficiently specific anchor gene (Donoho and Stodden, 2003). We therefore obtain the following identifiability result.

**Theorem 1.** *For the stGP model, assume that Conditions* (C1)-(C2) *hold, and at least one of* (C3a) *or* (C3b) *is satisfied. Then the parameter* $(W, C, D, \sigma_e^2)$ *is identifiable from the marginal distribution of* $Y$*, up to a permutation of the program labels.*

*Proof.* Suppose that two parameter tuples $(W, C, D, \sigma_e^2)$ and $(\widetilde{W}, \widetilde{C}, \widetilde{D}, \widetilde{\sigma}_e^2)$ generate the same marginal covariance matrix $\Omega$. Write $M_A = W^\top CW \in \mathbb{R}_+^{G\times G}$ and $M_S = W^\top DW \in \mathbb{R}_+^{G\times G}$, and define $\widetilde{M}_A$ and $\widetilde{M}_S$ analogously. Equating the two covariance representations gives

$$M_A \otimes K_A + M_S \otimes K_S + \sigma_e^2 I_{NG} = \widetilde{M}_A \otimes K_A + \widetilde{M}_S \otimes K_S + \widetilde{\sigma}_e^2 I_{NG}.$$

By Condition (C1), we then have $M_A = \widetilde{M}_A$, $M_S = \widetilde{M}_S$, and $\sigma_e^2 = \widetilde{\sigma}_e^2$. Therefore, it remains to show the identifiability of $(W, C, D)$ from $(M_A, M_S)$.

Part 1. We first consider the case in which Condition (C3a) holds. Since some programs may be purely temporal or purely spatial, it is more convenient to work with the total variance matrix $M = M_A + M_S = W^\top (C + D) W \in \mathbb{R}_+^{G \times G}$. Subsequently, define $V = (C + D)^{1/2} W \in \mathbb{R}_+^{p \times G}$ and $\widetilde{V} = (\widetilde{C} + \widetilde{D})^{1/2} \widetilde{W} \in \mathbb{R}_+^{p \times G}$. Let $\Gamma = (C + D)^{-1/2} C (C + D)^{-1/2} = \mathrm{diag}(\gamma_1, \cdots, \gamma_p)$ with $\widetilde{\Gamma}$ defined similarly. By Condition (C2), each diagonal entry of $C + D$ is strictly positive, and $W$ has full row rank $p$. Hence $V$ also has full row rank $p$. Moreover, we have

$$V^\top V = M = \widetilde{M} = \widetilde{V}^\top \widetilde{V}.$$

Therefore, there should exist an orthogonal matrix $R \in \mathbb{R}^{p \times p}$ such that $\widetilde{V} = RV$. Since

$$M_A = V^\top \Gamma V = \widetilde{V}^\top \widetilde{\Gamma} \widetilde{V} = V^\top R^\top \widetilde{\Gamma} R V,$$

and the fact that $V$ has full row rank, multiplying by a left inverse of $V$ on both sides leads to $\Gamma = R^\top \widetilde{\Gamma} R$. Under Condition (C3a), the diagonal entries of $\Gamma$ and $\widetilde{\Gamma}$ are pairwise distinct. This implies that the only orthogonal matrix that conjugates $\widetilde{\Gamma}$ to $\Gamma$ is the signed permutation matrix, i.e., $R = SP$, where $P$ is a permutation matrix and $S$ is diagonal with entries in $\{\pm 1\}$. This makes $\widetilde{\Gamma}$ and $\Gamma$ have exactly the same diagonal entries, only with different orders.

Moreover, recall that both $V$ and $\widetilde{V} = RV$ are entrywise nonnegative, negative sign changes are impossible. Therefore, we have $S = I_p$, so $R = P$ is an ordinary permutation matrix. It follows then $\widetilde{V} = PV$. Let $v_j^\top$ denote the $j$-th row of $V$. Since each row of $W$ lies on the simplex, we then have

$$v_j^\top 1_G = (\sigma_j^2 + \tau_j^2)^{1/2} w_j^\top 1_G = (\sigma_j^2 + \tau_j^2)^{1/2}.$$

Therefore, for each program $j$, we may recover

$$w_j = \frac{v_j}{v_j^\top 1_G}, \qquad \sigma_j^2 = \gamma_j (v_j^\top 1_G)^2, \qquad \tau_j^2 = (1 - \gamma_j)(v_j^\top 1_G)^2.$$

Thus $(W, C, D)$ is uniquely determined up to a permutation of the program labels. This proves the identifiability under Condition (C3a).

PART 2. We next consider the case in which Condition (C3b) holds. Fix a program $j$ and let $g_j$ be its anchor gene. Since $w_{kg_j} = 0$ for all $k \neq j$, we have

$$M_{g_j x} = \sum_{\ell=1}^{p} (\sigma_\ell^2 + \tau_\ell^2) w_{\ell g_j} w_{\ell x} = (\sigma_j^2 + \tau_j^2) w_{j g_j} w_{jx}$$

for $x = 1, \cdots, G$. Summing over $x$ and using $1_G^\top w_j = 1$, we then have $\sum_{x=1}^{G} M_{g_j x} = (\sigma_j^2 + \tau_j^2) w_{j g_j}$. Therefore, the entire row $w_j$ is identified by $w_{jx} = M_{g_j x} / \sum_{g'=1}^{G} M_{g_j g'}$ for $x = 1, \cdots, G$. Once $w_j$ is identified, we can recover the variance components from the original matrices $M_S$ and $M_A$. Recall that $(M_A)_{g_j x} = \sigma_j^2 w_{j g_j} w_{jx}$ and $(M_S)_{g_j x} = \tau_j^2 w_{j g_j} w_{jx}$, we then have $\sigma_j^2 = \sum_{g'=1}^{G} (M_A)_{g_j g'} / w_{j g_j}$ and $\tau_j^2 = \sum_{g'=1}^{G} (M_S)_{g_j g'} / w_{j g_j}$. Applying this argument to each program $j = 1, \cdots, p$, we can uniquely identify $(W, C, D)$, up to a permutation of the program labels. This proves the identifiability under Condition (C3b). □

Theorem 1 shows that the stGP model is identifiable on a biologically meaningful restricted parameter class. This result also clarifies how the simplex constraint and the covariance structure play complementary roles. The simplex constraint fixes the row scale of $W$, while either variance heterogeneity across programs or anchor-gene separability resolves the remaining rotational ambiguity. In particular, Condition (C3a) distinguishes programs through their temporal-to-total variance proportions, whereas Condition (C3b) distinguishes programs through sufficiently specific marker genes.

#### A.6 Implementation Details

We next describe the construction of the kernel matrices in the Gaussian processes. More specifically, let $A_1, \cdots, A_T$ denote the standardized chronological ages of the $T$ individuals. For the temporal dependence, we consider two choices, i.e., the Gaussian kernel and the autoregressive (AR) kernel. For the Gaussian kernel, we define $K_{\text{age}}(t, t') = \exp\big\{-(A_t - A_{t'})^2/\gamma_{\text{age}}\big\}$, where the bandwidth is given by $\gamma_{\text{age}} = d_{\text{age}}^2/c$, $d_{\text{age}} = \text{median}\big\{|A_t - A_{t'}| : 1 \le t < t' \le T\big\}$ is calculated by the median difference technique (Shang and Zhou, 2022) and $c$ is some constant. Alternatively, we can optionally consider an AR kernel as $K_{\text{age}}(t, t') = \rho^{|r(A_t) - r(A_{t'})|}$, where $r(\cdot)$ denotes the rank of an age value among the observed ages and the correlation depends on the ordering of individuals rather than on the raw age values. This is more realistic when ages are sparse or available only as ordered stages. Next, for the spatial dependence, define $S_i^t \in \mathbb{R}^2$ as the standardized spatial coordinate of cell $i$ for each individual $t$. Then, we construct the spatial kernel matrix for individual $t$ as $K_{\text{spa}}^t(i, i') = \exp\big\{-\|S_i^t - S_{i'}^t\|^2/\gamma_{\text{spa}}\big\}$, where the bandwidth is given by $\gamma_{\text{spa}} = \text{median}\big\{d_{\text{spa}}^{(1)}, \cdots, d_{\text{spa}}^{(T)}\big\}^2/c$, and $d_{\text{spa}}^{(t)} = \text{median}\big\{\|S_i^t - S_{i'}^t\| : 1 \le i < i' \le N_t,\ \|S_i^t - S_{i'}^t\| > 0\big\}$ refers to the per-slice median values.

For the multi-program case with $p > 1$, we fit the stGP model using a backfitting procedure with automated rank selection (Wang et al., 2024). Specifically, we first choose an upper bound $p_{\max} = 10$ for the number of programs, and sequentially extract rank-1 stGP components from the residual. If $R_{j-1}$ denotes the residual before adding the $j$-th factor, then after fitting a candidate component $(h_j, w_j)$, we define the updated residual as $R_j = R_{j-1} - h_j w_j^\top$. The greedy addition stops when the relative incremental improvement $\delta_j = \big(\|R_{j-1}\|_F^2 - \|R_j\|_F^2\big)/\|R_{j-1}\|_F^2$ becomes smaller than 0.05 by default. Starting from this initialization, we cycle through factors $j = 1, \cdots, \widehat{p}_{\text{greedy}}$ and update one component at a time while holding the others fixed. The backfitting iterations are terminated when the relative changes in both the loading matrix $W$ and the variance components $(\sigma_j^2, \tau_j^2)$ fall below $10^{-4}$. In addition, since multi-factor decompositions might possibly yield highly similar loading vectors, we should monitor the pairwise similarities $\cos(w_j, w_{j'}) = w_j^\top w_{j'}/(\|w_j\| \, \|w_{j'}\|)$ for different program $j \neq j'$s. If the largest cosine similarity exceeds 0.9, we treat the corresponding pair as near-duplicate factors, remove them from the current fit, and sequentially re-extract rank-1 components from the resulting residual. After backfitting, each factor is further evaluated by its explained-energy fraction $\pi_j = \|h_j\|^2 \, \|w_j\|^2/\|Y\|_F^2$. Factors with $\pi_j < 0.01$ are removed, and the model is refit until no additional factor is discarded.

### B Supplementary Notes

#### B.1 Benchmarking Methods

We compared stGP with six baseline methods spanning non-spatial matrix factorization, spatial factorization and spatiotemporal factor models: PCA, NMF, SpatialPCA, MEFISTO, STAMP and Popari. For each analysis, the number of components, factors, topics or metagenes was set to the target rank used for the corresponding simulation or real-data benchmark.

**PCA.** We applied PCA using the scikit-learn v1.8.0 implementation. PCA was applied to the stacked log-scale expression matrix after gene centering. In real data benchmarks, counts were first normalized to a target sum of 250 per cell, transformed by `log1p`, and was fitted by singular value decomposition.

**NMF.** We applied NMF using the scikit-learn v1.8.0 implementation. NMF was applied to non-negative log-normalized expression, obtained by normalized to a target sum of 250 per cell followed by `log1p`. We used `init="nndsvda"`, with `max_iter=500` in the real data analysis.

**SpatialPCA.** We applied SpatialPCA v1.3.0 in R v4.5.3. SpatialPCA was fitted jointly across samples using per-section expression matrices and spatial coordinates, with a Gaussian block-diagonal spatial kernel. The scripts skipped SPARK-based spatially variable gene selection because the MERFISH panels were fixed, used the common gene intersection across samples, performed per-sample Seurat `LogNormalize` preprocessing and integrated samples by Seurat rPCA with dimensions 1–30, `k.anchor=5` and `k.filter=200`.

**MEFISTO.** We applied MEFISTO using `mofapy2` v0.7.3 and `mofax` v0.3.7. MEFISTO was fitted as a Gaussian factor model with a sparse Gaussian-process prior over covariates and the requested number of factors. We used `use_float32=True`, 1,000 inducing points, 1,000 training iterations, `convergence_mode="fast"`, `start_opt=10`, `opt_freq=10`, `scale_cov=False` and `model_groups=False`. The inducing-point fraction was computed as `n_inducing/n_cells` and clipped to the interval [`1e-4`, 0.8].

**STAMP.** We applied STAMP through `sctm` v0.1.3. Spatial neighbors were computed with Squidpy using $\max\{5, \mathrm{round}(0.006\, n_{\mathrm{cells}})\}$ neighbors, and STAMP was run with `mode="sgc"`, `sampler="W"`, `max_epochs=400`, `min_epochs=100`, learning rate 0.01 and batch size 256. Cell-topic proportions were used as cell embeddings, and topic-gene features were used as loadings where available. Since invalid parameter values might appear during training, a small schedule of gene cutoffs was tried until training produced non-NaN outputs.

**Popari.** We applied Popari v0.0.72 with `torch` v2.9.1. Popari was fitted as a multi-sample spatial metagene model on per-sample datasets. In real data benchmarks, counts were normalized to a target sum of 250 and transformed by `log1p`; up to 200 highly variable genes were selected from the merged normalized data before splitting into per-`mouse_id` Popari datasets. The real-data Popari runner used $\lambda_{\Sigma_x^{-1}} = 10^{-4}$ and $\lambda_{\bar{\Sigma}} = 10^{-4}$, the differential-lookup spatial-affinity mode, 10 NMF-style warm-up iterations and 200 main training iterations.

#### B.2 Simulation Design

The simulation studies evaluated whether stGP recovered three latent quantities unavailable in real data: sparse gene-program loadings, cell-level program embeddings, and their decomposition into temporal and within-section spatial effects. Each simulation study used 50 independent replicates.

**Latent spatiotemporal signal.** For section $t = 1, \dots, T$, the latent log-scale expression matrix $Y^t_{\log} \in \mathbb{R}^{N_t \times G}$ was generated as

$$Y^t_{\log} = H^t W + E^t, \qquad E^t_{ig} \sim N(0, \sigma_e^2),$$

where $W \in \mathbb{R}^{p \times G}$ contains simplex-normalized sparse gene-program loadings. The embedding for program $j$ was $h^t_{ij} = \alpha_j(t) + b^t_{ij}$, where $\alpha_j \sim N(0, \sigma^2_{j,\text{age}} K_{\text{age}})$ and $b^t_j \sim N(0, \tau^2_{j,\text{spa}} K^t_{\text{spa}})$. Thus, $K_{\text{age}}$ captured covariance across sections and $K^t_{\text{spa}}$ captured within-section spatial covariance. Ages were placed on a 3-30 grid, jittered by 15% of the grid spacing, and standardized before temporal kernel construction. Spatial coordinates were generated independently within each section and standardized. Sparse programs were generated by selecting $k$ active genes and drawing their weights from a Dirichlet distribution. The simulation used $T = 20$ sections, $G = 100$ genes, and $p = 4$ programs. Each section contained a random number of cells with $N_t = \max\{100, \text{round}(Z_t)\}$ and $Z_t \sim N(150, 15^2)$. Temporal and spatial covariances used Gaussian RBF kernels, $K_{\text{age}}(t, t') = \exp[-(A_t - A_{t'})^2/\gamma_{\text{age}}]$ and $K^t_{\text{spa}}(i, i') = \exp[-\|S^t_i - S^t_{i'}\|^2/\gamma_{\text{spa}}]$, with $\gamma_{\text{age}} = 0.5$ and $\gamma_{\text{spa}} = 2.0$. The program-specific variance components were $(\sigma^2_{1,\text{age}}, \dots, \sigma^2_{4,\text{age}}) = (10, 5, 0, 8)$ and $(\tau^2_{1,\text{spa}}, \dots, \tau^2_{4,\text{spa}}) = (6, 10, 8, 0)$, respectively. This created two programs with both temporal and spatial variation, one spatial-only program, and one temporal-only program. To mimic count data, marker genes, housekeeping genes, and mostly inactive genes were assigned different baseline-expression ranges, and each cell received a log-scale capture-efficiency offset. Counts were drawn from

$$\log \mu^t_{ig} = a_g + c^t_i + (H^t W)_{ig},$$

where $a_g$ is the gene baseline and $c^t_i$ is the cell-level offset calibrated to an expected library size near 500 counts. Observed counts were sampled as $Y^{t,\text{count}}_{ig} \sim \text{Poisson}(\mu^t_{ig})$

and used for method fitting.

Estimated programs were matched to ground truth by the Hungarian algorithm, using absolute loading values, top-10% sparsification, and row-wise simplex projection to compare signed and non-negative methods. Loading recovery was measured by the mean Hellinger score, $1 - d_H$, and the top-$k$ hit rate for active genes. Embedding recovery was measured by the mean Pearson correlation between matched stacked embeddings. Additional metrics, including total-variation loading similarity, 90%-mass gene-set F1 score, Spearman embedding correlation, and wall-clock runtime, are reported in Supplementary Figure S5.

Additional simulations tested rank selection, kernel misspecification, Gaussian log-scale observations, and a single-cell-like setting. For rank selection, Gaussian log-scale data with $p_{\text{true}} = 4$ were fitted with $p_{\max} = 10$; stGP selected $\widehat{p} = 4$ in all 50 replicates after extracting, pruning, and merging candidate components (Supplementary Figure S1). Kernel sensitivity was assessed by fitting an AR(1) temporal kernel to RBF-generated data, varying the fitted AR(1) correlation for AR(1)-generated data, perturbing the spatial bandwidth, and evaluating automatic bandwidth selection (Supplementary Figure S2). The Gaussian benchmark reused the four-program design without count generation (Supplementary Figure S3). The single-cell-like benchmark set spatial covariance to zero and compared stGP, PCA, and NMF on Poisson counts (Supplementary Figure S4).

### C Supplementary Figures

#### C.1 Simulations

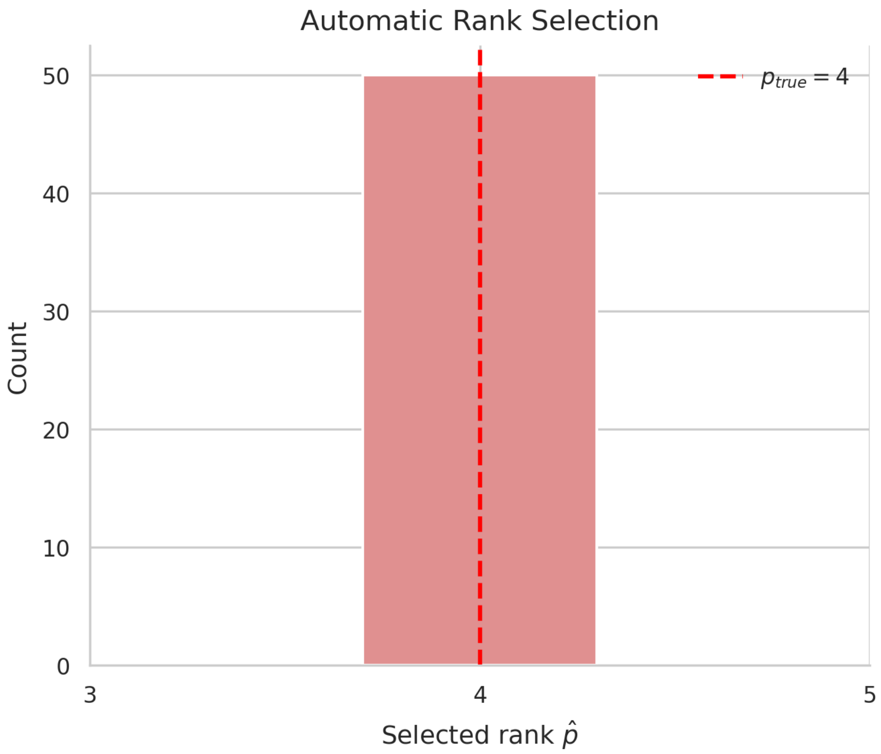

Supplementary Figure S1: **Automatic rank selection recovers the true number of programs.** The automatic stGP backfitting procedure was run with $p_{\max} = 10$ in simulated datasets with $p_{\text{true}} = 4$. The histogram shows the selected rank $\widehat{p}$ across 50 replicates. The dashed red line marks the true rank.

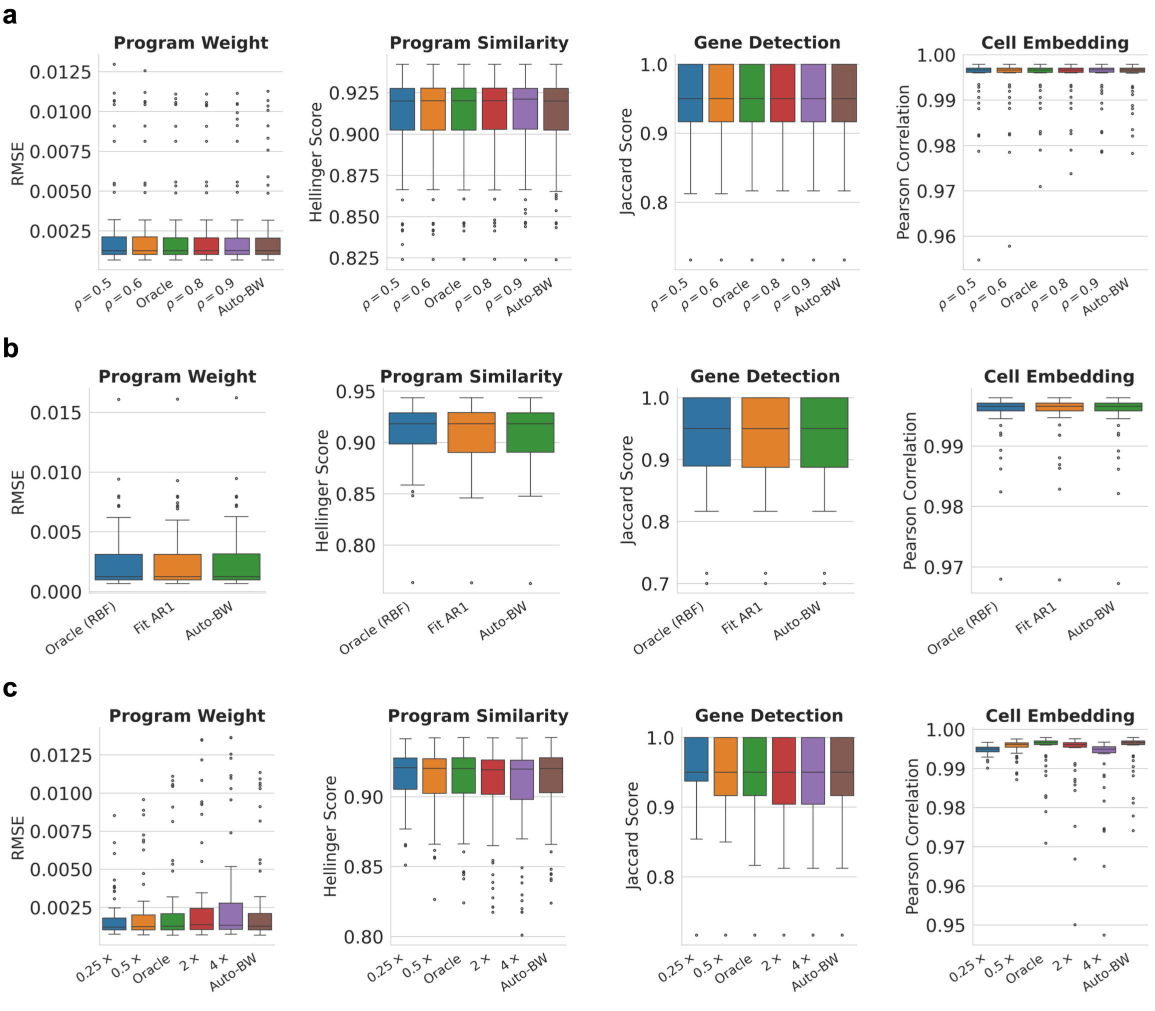

Supplementary Figure S2: **stGP is robust to temporal and spatial kernel misspecifications. a,** Recovery metrics across AR(1) temporal kernels and automatic bandwidth selection when the data are generated from an AR(1) kernel with $\rho = 0.7$. **b,** Recovery metrics when fitting an AR(1) temporal kernel to data generated from a Gaussian temporal kernel, compared with the oracle fit and automatic bandwidth selection. **c,** Recovery across spatial bandwidth multipliers relative to the oracle setting. Boxplots summarize program-weight RMSE, program similarity, gene detection and cell-embedding correlation across simulation replicates.

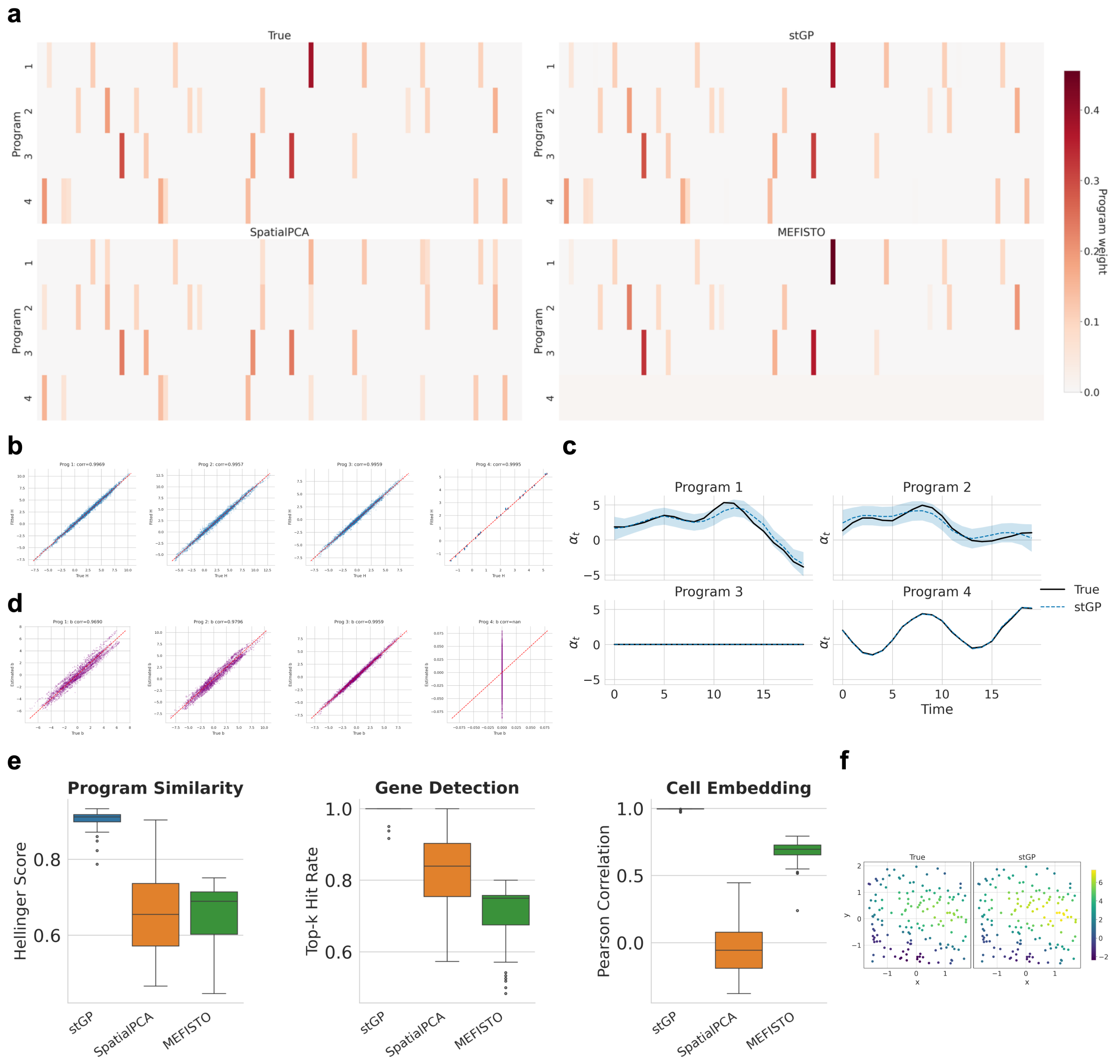

Supplementary Figure S3: **stGP accurately recovers gene programs in log-scale simulations.** **a,** True and estimated gene loadings for stGP, SpatialPCA and MEFISTO in one replicate. **b,** Pearson correlation between estimated and true cell embeddings for each program. **c,** Recovered temporal effects compared with the ground truth. **d,** Pearson correlation between estimated and true within-section spatial effects. **e,** Quantitative recovery metrics for gene programs, active genes and cell embeddings across 50 replicates. **f,** True and stGP-recovered spatial activity for a representative program.

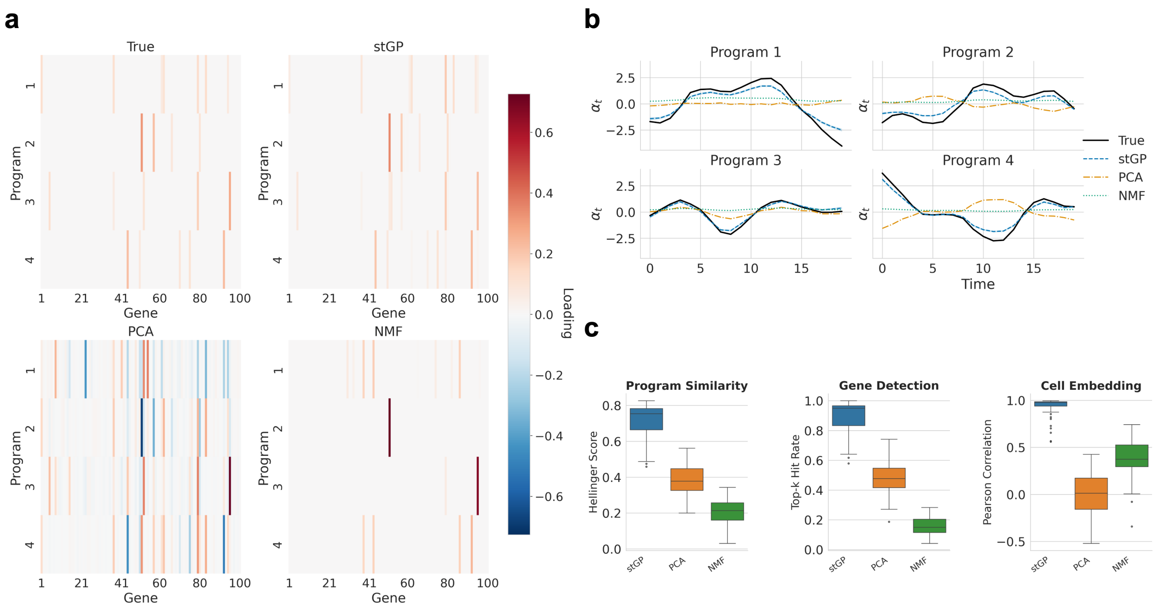

Supplementary Figure S4: **stGP reduces to a temporal model when spatial effects are absent for single-cell simulations.** The simulation generated Poisson count data with temporal gene programs while setting the spatial variance components to zero. **a,** True and estimated gene loadings for stGP, PCA and NMF in a representative replicate. **b,** Recovered temporal effects compared with the ground truth. **c,** Quantitative recovery metrics of gene programs and cell embeddings across 50 replicates.

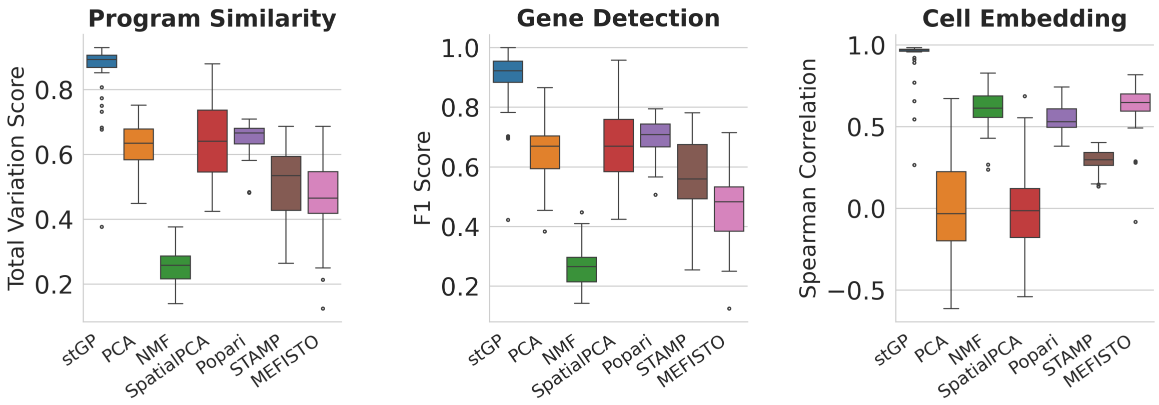

Supplementary Figure S5: **Additional simulation metrics support the main recovery benchmark.** Program recovery, gene detection and cell embedding recovery were evaluated using total variation, F1 score and Spearman correlation, respectively. Across these complementary metrics, stGP retained the strongest overall recovery among the compared methods.

#### C.2 Human Aging DLPFC

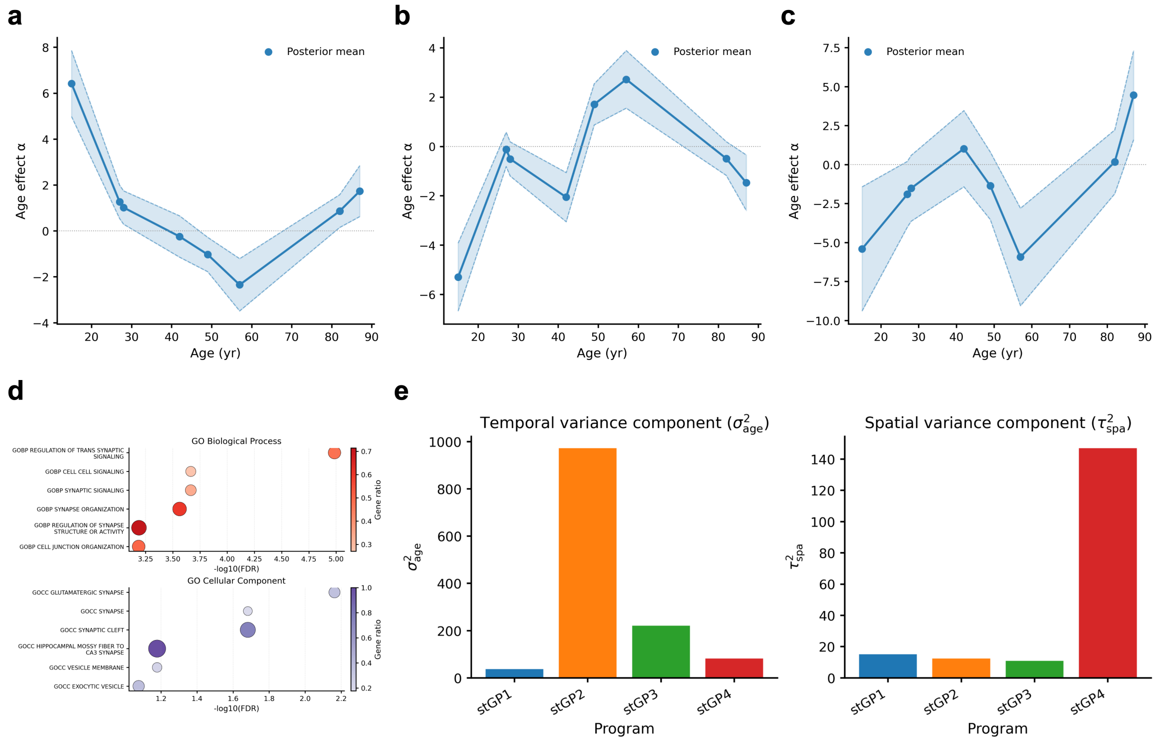

Supplementary Figure S6: **stGP resolves temporally and spatially structured excitatory-neuron programs in human DLPFC sections. a-c,** Estimated temporal effects of stGP1 (a), stGP2 (b) and stGP4 (c) for excitatory-neuron programs across donor age. **d,** Gene Ontology enrichments for stGP1, showing synaptic and glutamatergic terms consistent with excitatory-neuron identity. **e,** Estimated temporal and spatial variance components for the four excitatory-neuron stGP programs. stGP1-stGP3 capture layer-associated spatiotemporal structure, whereas stGP4 is dominated by within-section spatial variation.

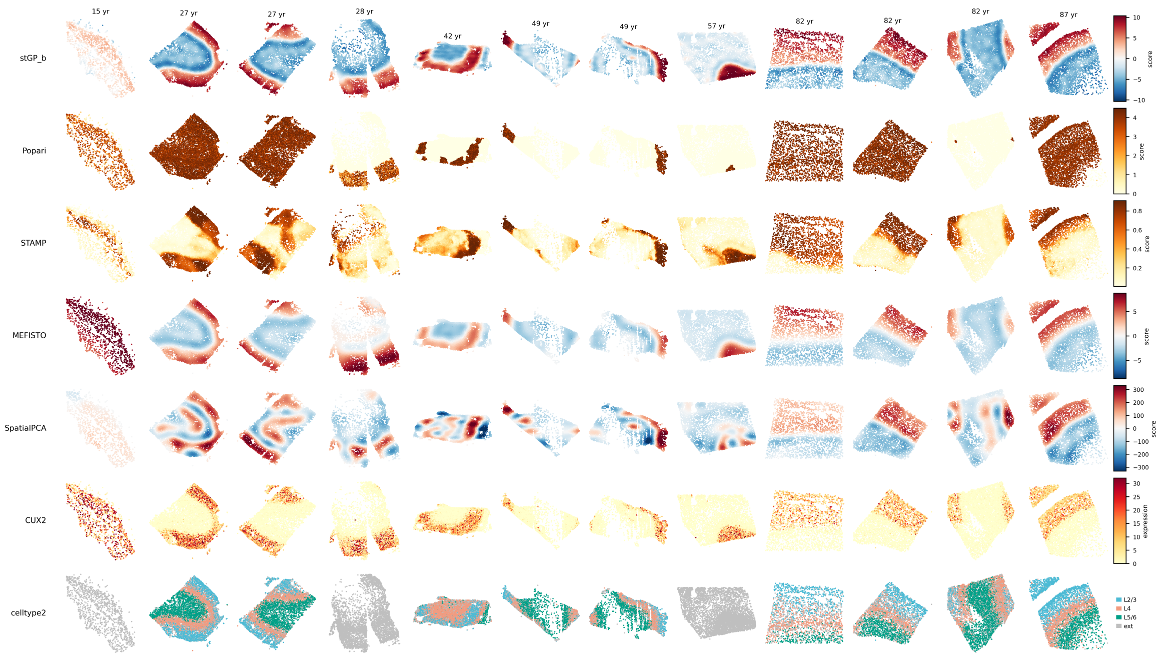

Supplementary Figure S7: **Benchmarking recovery of the L2/3-associated excitatory-neuron pattern across unaligned human DLPFC sections.** Spatial embeddings or domains from stGP and baseline methods are shown across the 12 human DLPFC MERFISH sections and compared with *CUX2* expression and reference excitatory-neuron subtype annotations. stGP spatial activity recovers a continuous upper-layer band across sections with different ages and orientations, whereas several baseline embeddings are fragmented, diffuse or dominated by section-specific structure.

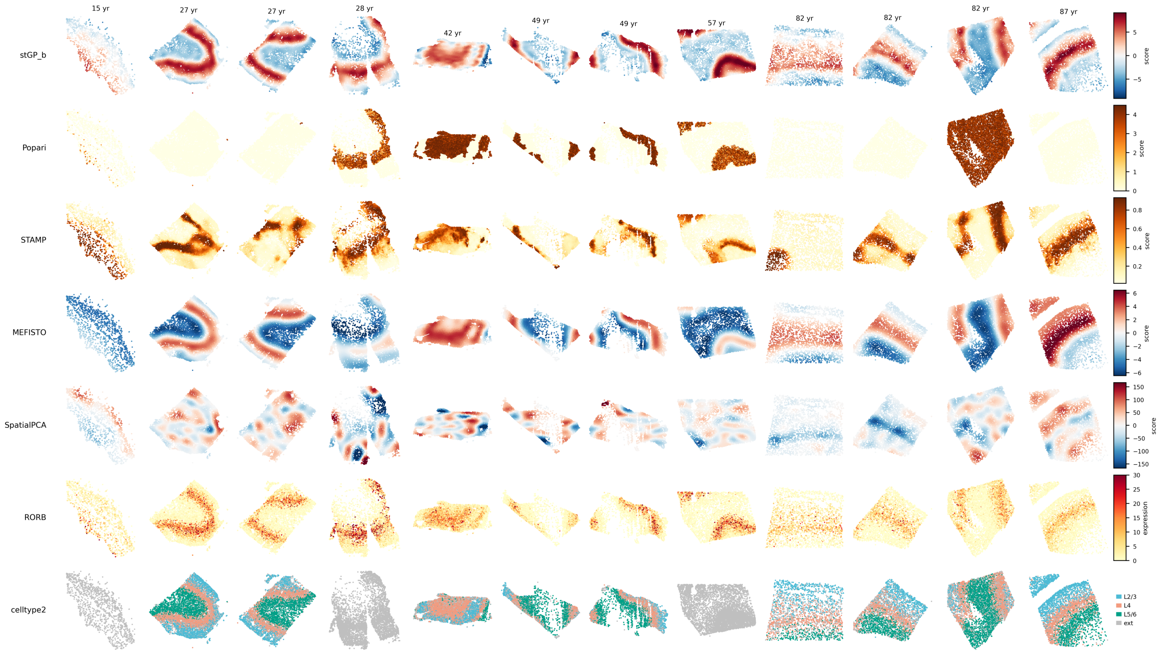

Supplementary Figure S8: **Benchmarking recovery of the L4-associated excitatory-neuron pattern across unaligned human DLPFC sections.** Spatial embeddings or domains from stGP and baseline methods are shown across the 12 human DLPFC MERFISH sections and compared with *RORB* expression and reference L4 excitatory-neuron subtype annotations. stGP recovers a narrow middle-layer band that is consistent across sections, supporting the main-text comparison of layer-marker correlations and domain-recovery metrics.

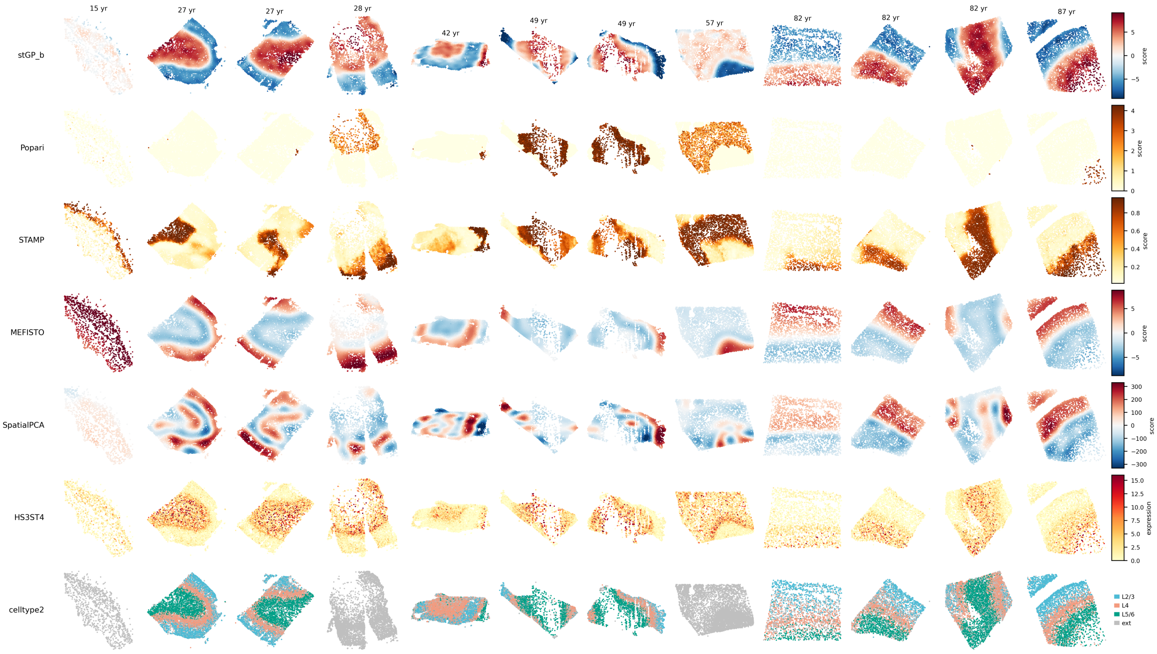

Supplementary Figure S9: **Benchmarking recovery of the L5/6-associated excitatory-neuron pattern across unaligned human DLPFC sections.** Spatial embeddings or domains from stGP and baseline methods are shown across the 12 human DLPFC MERFISH sections and compared with *HS3ST4* expression and reference L5/6 excitatory-neuron subtype annotations. stGP captures deep-layer spatial organization across sections, while baseline methods show weaker continuity, stronger patchiness or less precise boundaries in several sections.

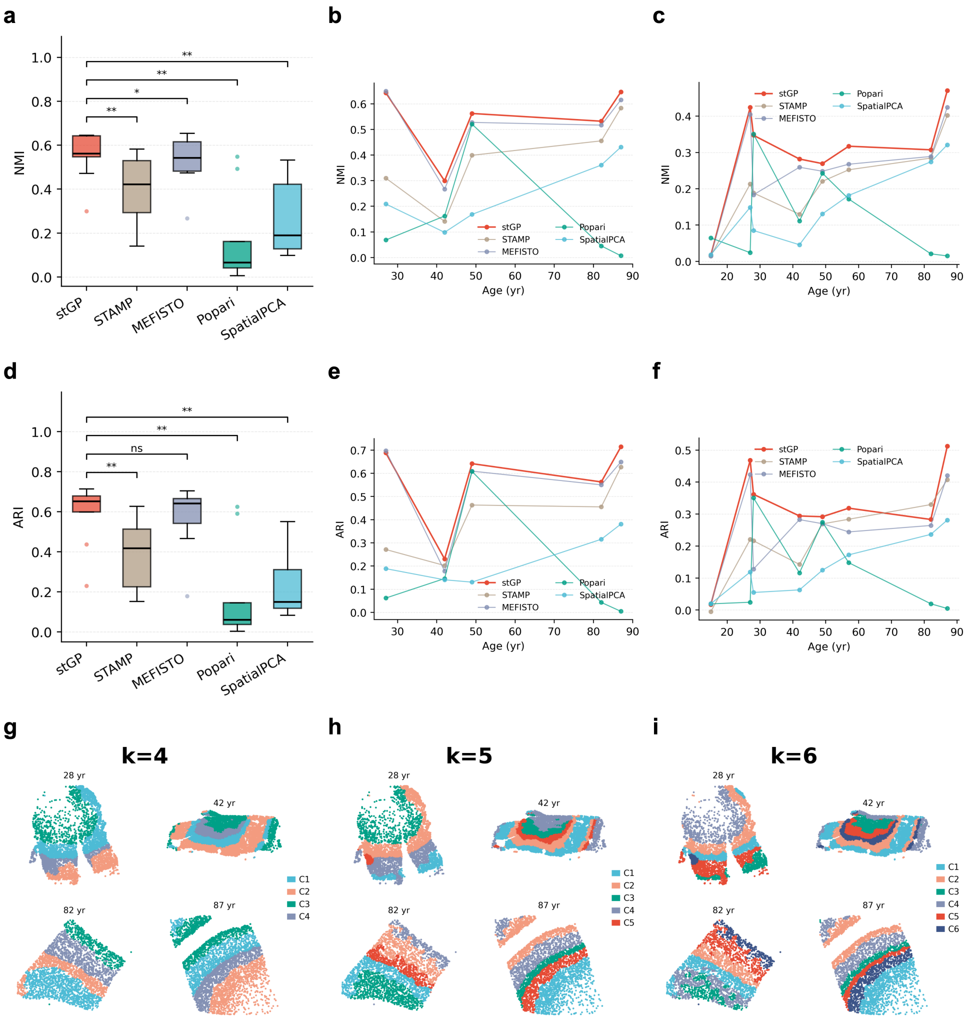

Supplementary Figure S10: **stGP spatial embeddings support robust laminar clustering and finer sublaminar structure. a,d,** Adjusted rand index (ARI) and normalized mutual information (NMI) for clustered spatial domains versus cell subtypes in slices with accurate annotations. **b,e,** Per-age summaries of NMI (b) and ARI (e) for clustered spatial domains versus cell subtype annotations. **c,f,** Per-age summaries of NMI (c) and ARI (f) for clustered spatial domains versus marker-defined labels. **g-i,** Clustered spatial domains based on stGP spatial embeddings with increasing cluster numbers ($k = 4$, 5 and 6), demonstrating that the same embedding can resolve both major layers and additional sublaminar domains.

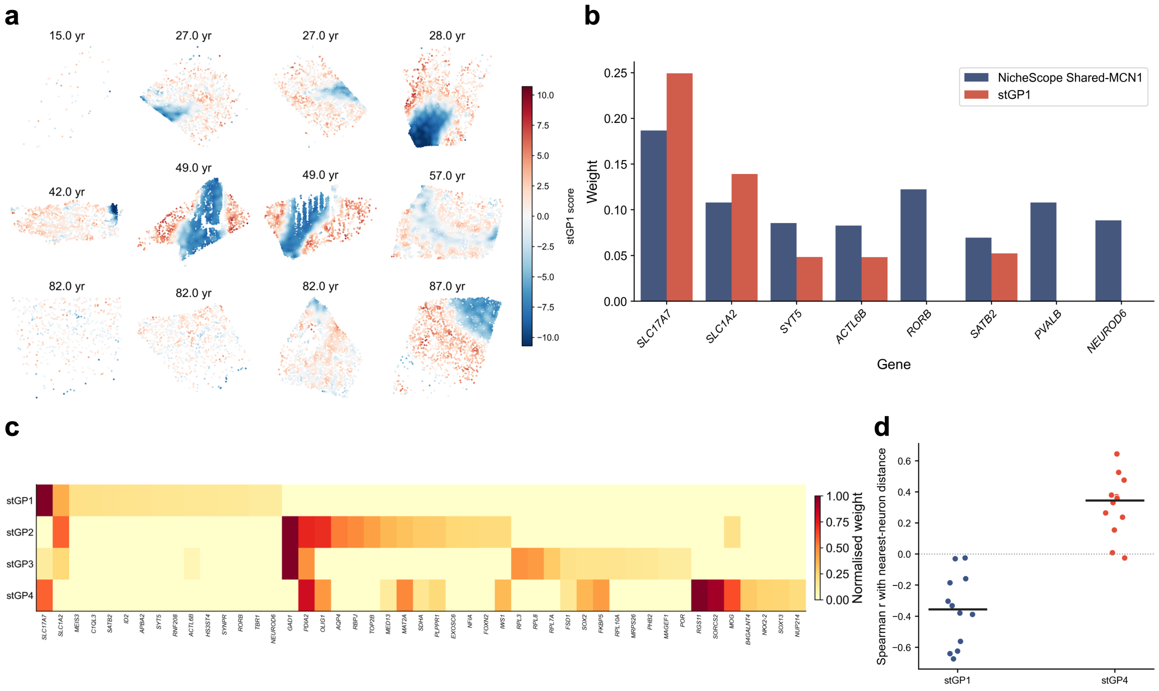

Supplementary Figure S11: **stGP fits in human aging DLPFC oligodendrocytes, as well as its comparison with NicheScope.** **a,** Spatial distribution of oligodendrocyte stGP1 activities across the 12 human DLPFC MERFISH sections. **b,** Gene loading comparison between NicheScope Shared-MCN1 and stGP1 for genes shared by their marker sets. **c,** Normalized gene-loading heatmap for stGP1-stGP4, using the union of top positive genes across programs. **d,** Section-level Spearman correlation between stGP1/stGP4 spatial embedding and distance to the nearest neuron. For each section, nearest-neuron distance was computed from all neuronal cells and assigned to each oligodendrocyte. Each point denotes one section, and black horizontal bars denote medians. Positive values indicate higher program scores farther from neuronal cell bodies, whereas negative values indicate higher scores nearer to neurons. stGP4 showed mostly positive correlations (median $\rho \approx 0.35$), whereas stGP1 showed negative correlations (median $\rho \approx -0.36$), arguing against a simple neuronal-proximity or neuronal-contamination explanation for stGP4 hotspots as a negative control.

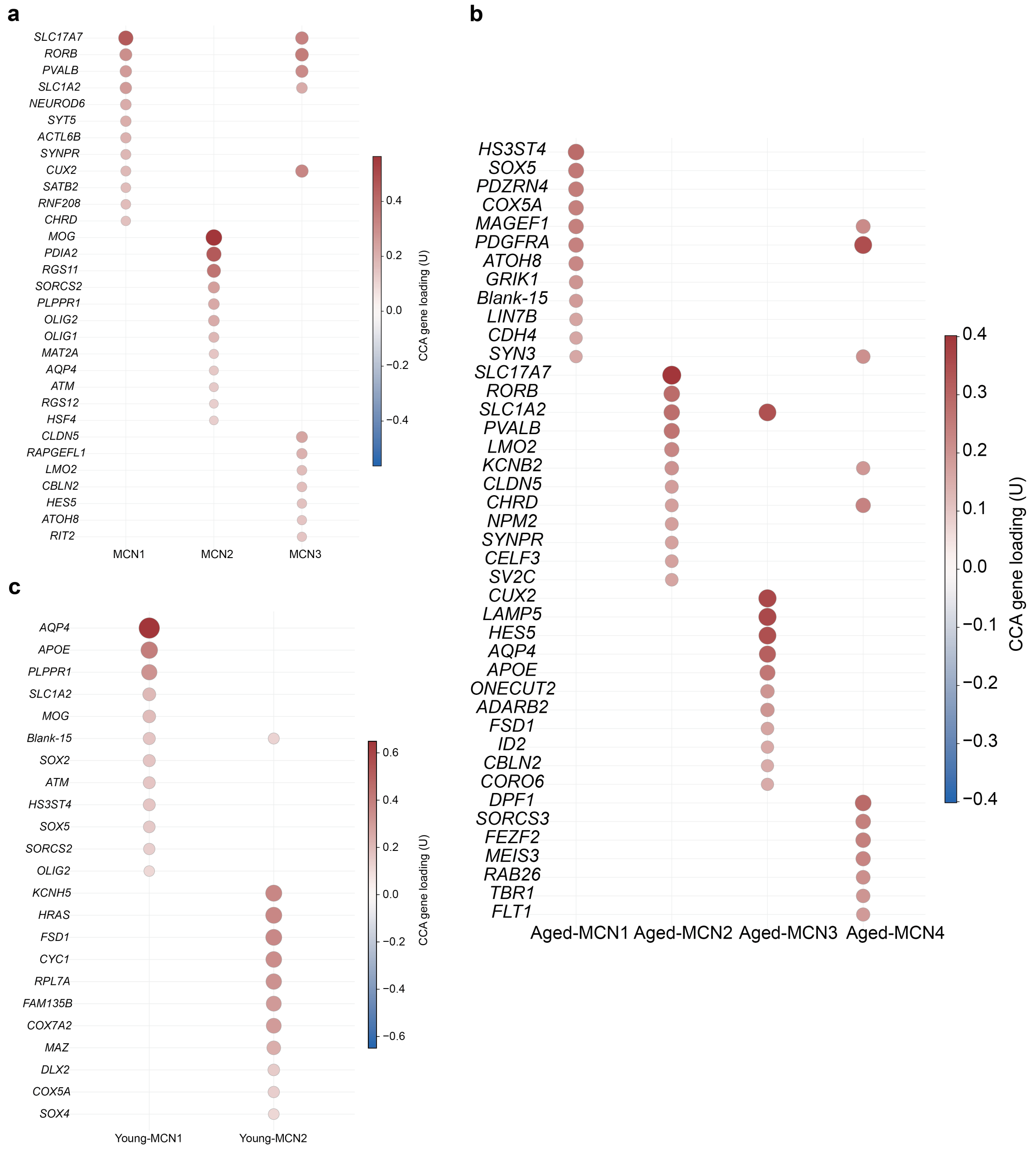

Supplementary Figure S12: **NicheScope CCA gene loadings for shared and age-specific human DLPFC multicellular niches.** Dot plots show canonical-correlation-analysis (CCA) gene loadings $U$ for NicheScope multicellular niches (MCNs). **a,** Shared MCN1-MCN3 gene-loading signatures estimated across young and aged sections. **b,** Aged-specific MCN1-MCN4 gene-loading signatures. **c,** Young-specific MCN1 and MCN2 gene-loading signatures.

#### C.3 Mouse Aging Brain

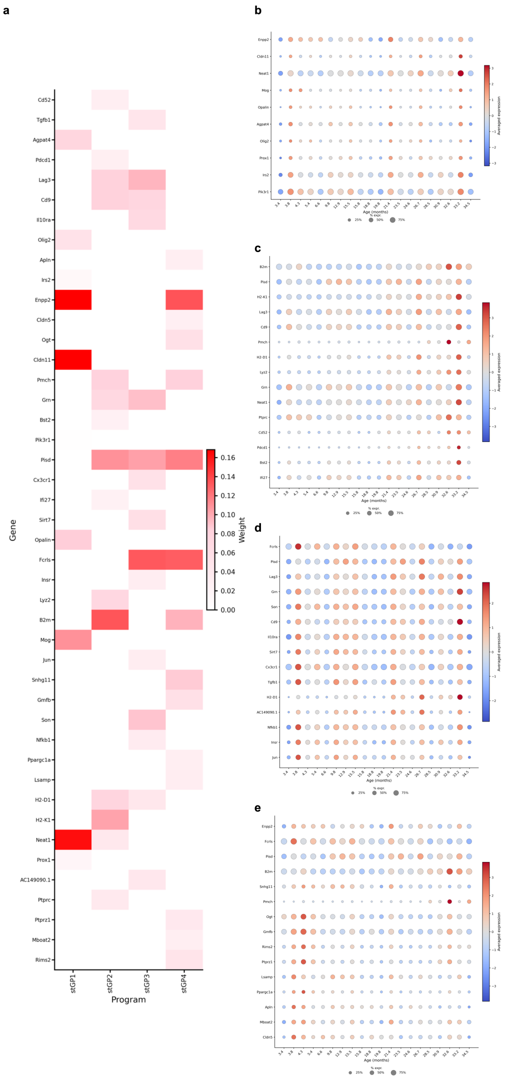

Supplementary Figure S13: **stGP resolves distinct gene programs in mouse aging brain microglia. a,** Gene-loading vectors for the four microglial stGP programs. **b-e,** Age-resolved average expression of genes for stGP1-stGP4. stGP1 (b) is enriched for myelin- and glial-associated genes including *Enpp2*, *Cldn11*, *Mog* and *Opalin*. stGP2 contains immune and major histocompatibility complex class I (MHC-I)-associated genes including *B2m*, *H2-K1* and *H2-D1*, and interferon-responsive genes including *Bst2* and *Ifi27*. stGP3 contains microglial homeostatic and regulatory genes including *Fcrls*, *Cx3cr1*, *Il10ra* and *Tgfb1*. stGP4 contains a mixed glial- and vascular-associated loading pattern including *Enpp2*, *Fcrls*, *Cldn5* and *Apln*.

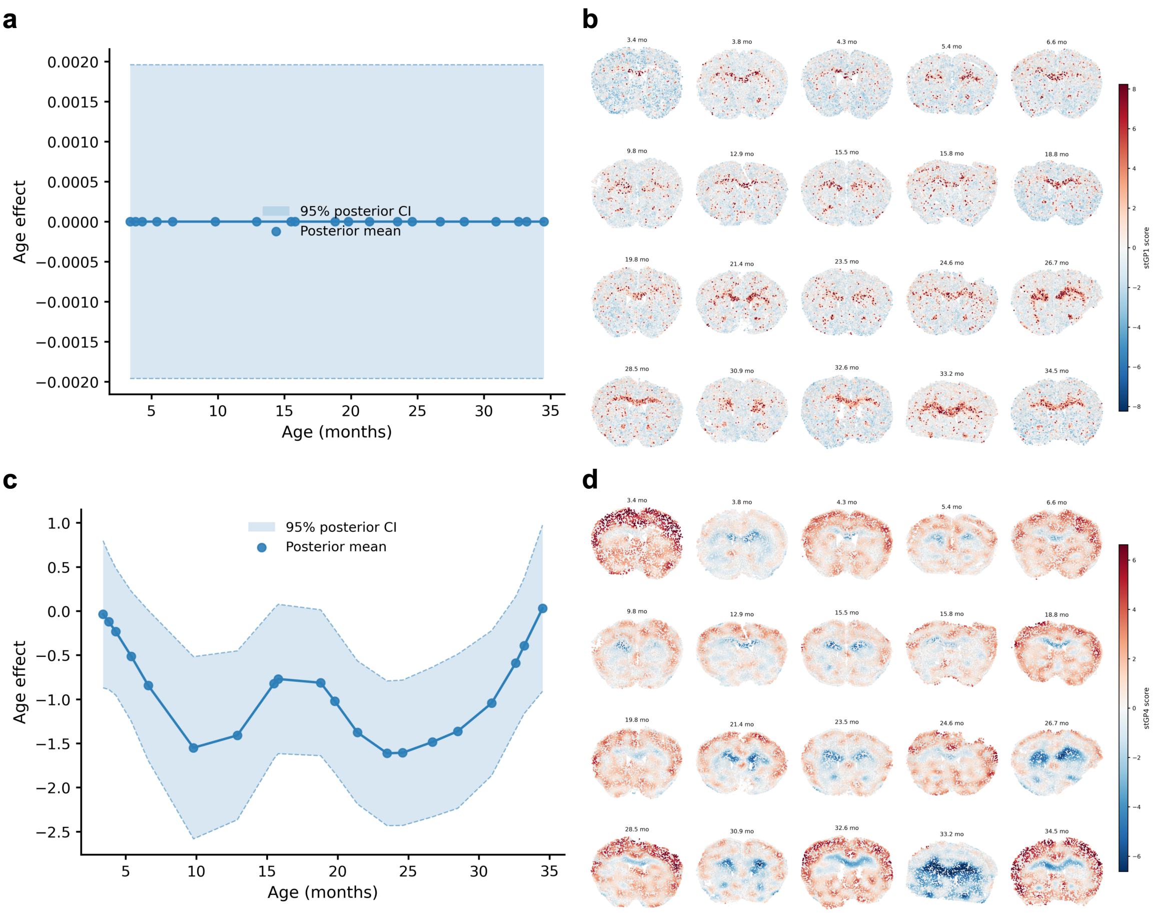

Supplementary Figure S14: **Temporal and spatial decomposition of the spatially patterned programs stGP1 and stGP4 in mouse aging brain microglia.** **a,** Temporal effect of stGP1. The posterior mean remains close to zero across the sampled age range, indicating little sample-level temporal shift for this program. **b,** Spatial activity maps for stGP1. High stGP1 activity is concentrated in the CC/ACO white-matter region across sections, consistent with a stable anatomical microglial program. **c,** Temporal effect of stGP4. stGP4 shows a nonlinear temporal trend with substantial posterior uncertainty, indicating limited evidence for a consistent sample-level temporal shift. **d,** Spatial activity maps for stGP4. stGP4 activity is mainly distributed in cortical regions.

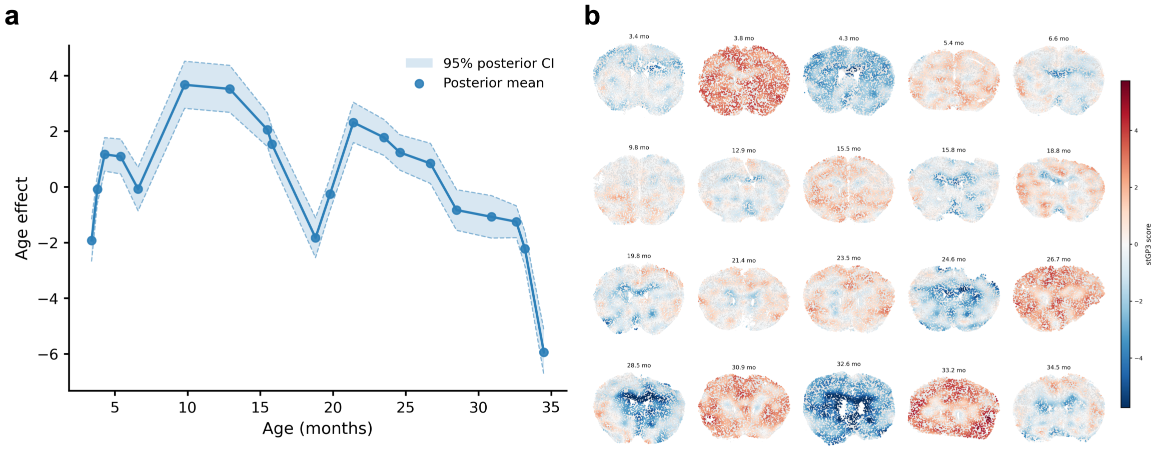

Supplementary Figure S15: **stGP3 captures a whole-brain declining program with microglial homeostatic and regulatory functions.** **a,** Temporal effect of stGP3. The temporal effect peaks in early to midlife and then declines, with the lowest posterior mean in the oldest section. **b,** Spatial activity maps for stGP3. stGP3 shows little spatial effect, as the estimated spatial variance component $\tau^2_{\text{spa}}$ is nearly zero. Instead, stGP3 activity is broadly distributed across brain sections, consistent with the homeostatic and regulatory gene program being present in most microglia and declining globally during late aging.

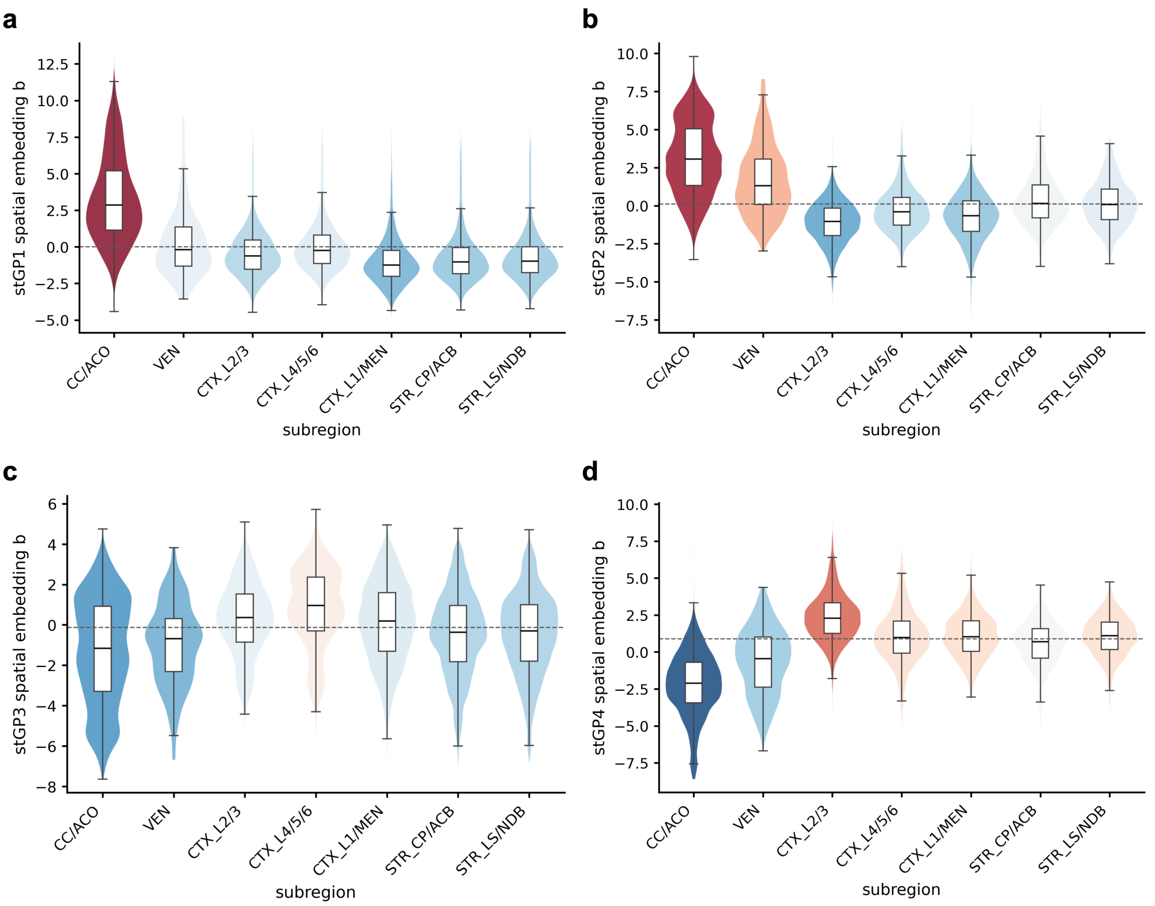

Supplementary Figure S16: **Subregion enrichment separates spatially localized and broadly distributed microglial stGP programs. a-d,** Distributions of the spatial activity $b$ for stGP1 (a), stGP2 (b), stGP3 (c) and stGP4 (d) across annotated mouse brain subregions. Violin plots show the cell-level distribution within each subregion, and boxplots summarize the median and interquartile range. stGP1 and stGP2 show the strongest positive spatial activity in the corpus callosum/anterior commissure (CC/ACO) region, while stGP2 also shows positive enrichment in the ventricle-adjacent region (VEN). stGP4 is enriched in cortex (especially CTX_L2/3) and depleted in CC/ACO. By contrast, stGP3 shows no dominant positive subregion enrichment, supporting its interpretation as a broadly distributed temporal program marked by late-life decline in homeostatic and regulatory gene activity.

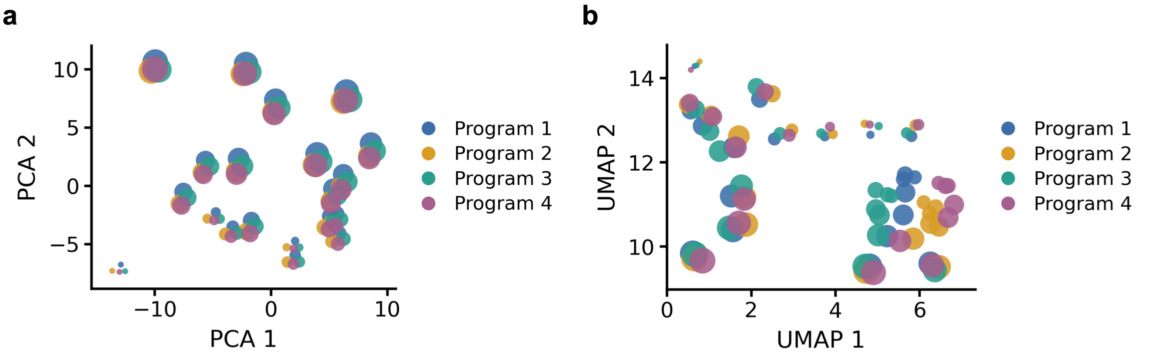

Supplementary Figure S17: **STAMP (Zhong et al., 2024) results for mouse aging brain microglia. a,** PCA embedding of the STAMP program representations. **b,** UMAP embedding of the same STAMP program representations. STAMP assumes fixed cell-level program activity across time and temporally smoothed latent programs. However, the PCA and UMAP embeddings suggest that the representations still cluster by sample rather than along an age trajectory. Thus, in this analysis, STAMP did not recover a clear temporal trajectory for the identified microglial programs.

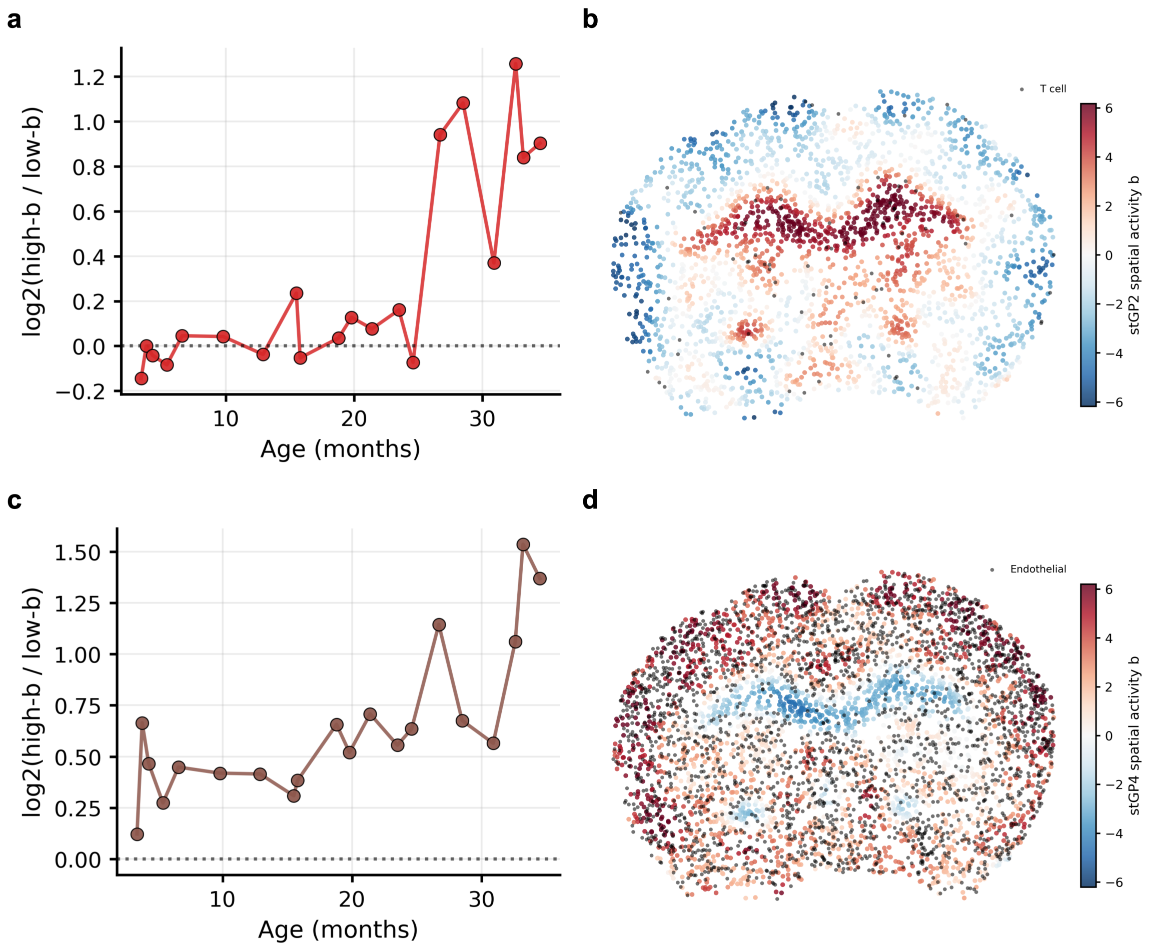

Supplementary Figure S18: **Spatially active microglial programs colocalize with T-cell and endothelial neighborhoods.** **a,** Age-resolved $\log_2$ ratio of T-cell abundance around microglia with high versus low stGP2 residual spatial activity $b$. High-$b$ and low-$b$ microglia were defined within each age slice as the top and bottom quartiles of the corresponding stGP residual spatial activity. Neighboring T cells or endothelial cells were counted in the 20-50 $\mu$m neighborhood shell around each microglial cell. **b,** Spatial overlay of stGP2 activity and T cells in the oldest brain section (34.5 months). **c,** Age-resolved $\log_2$ ratio of endothelial-cell abundance around high-$b$ versus low-$b$ microglia for stGP4. **d,** Spatial overlay of stGP4 activity and endothelial cells in the oldest brain section (34.5 months).

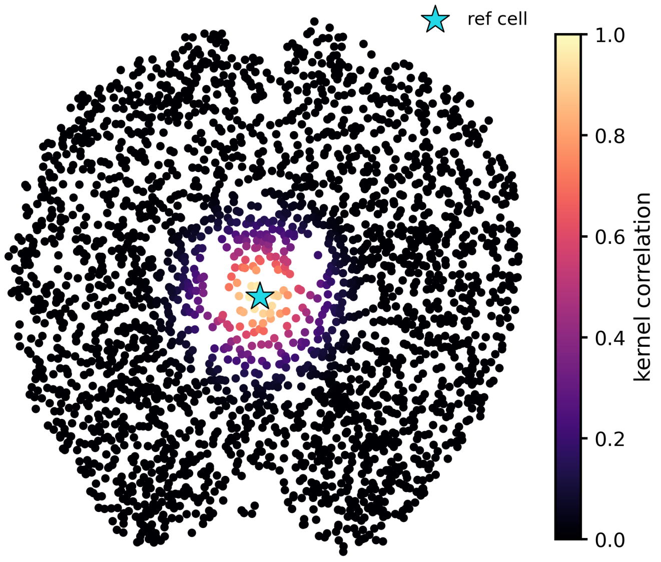

Supplementary Figure S19: **Reference-cell spatial correlation.** In the constructed spatial kernel, the cyan star marks the reference cell, and all other cells are colored by their spatial kernel correlation with this reference cell. The high-correlation neighborhood is concentrated around the reference cell and decays with spatial distance, illustrating the locality encoded by the constructed spatial kernel.

#### C.4 Mouse Injured Kidney

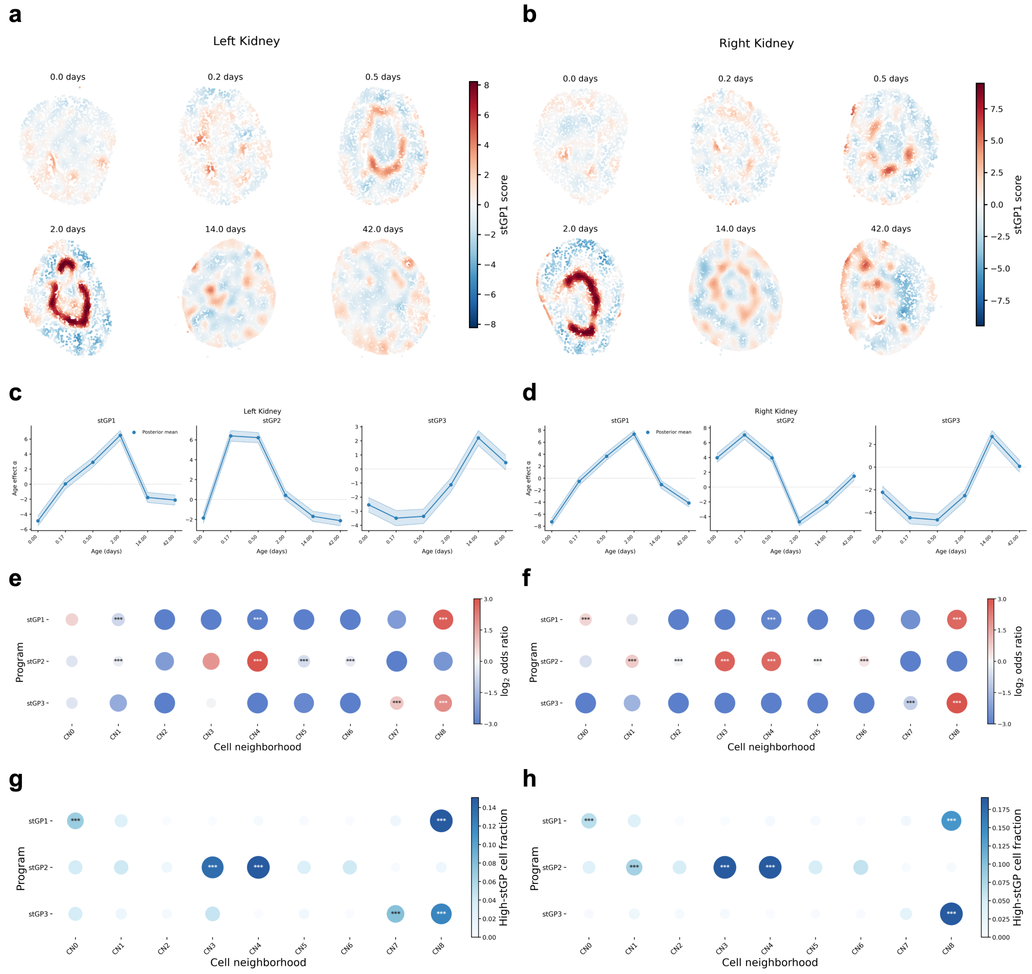

Supplementary Figure S20: **Including sham as a baseline preserves the stage-specific immune repair programs across bilateral kidneys. a,b,** Spatial activities of the matched sham-inclusive immune repair-transition program learned from left-kidney (a) and right-kidney (b) immune cells. The day-2 activation in restricted annular bands around the injury sites is preserved in both fits. **c,d,** Temporal effects for the three sham-inclusive immune programs in the left (c) and right (d) kidney fits. **e,f,** CN enrichment of high-scoring immune cells in the left (e) and right (f) kidney fits, shown as $\log_2$ odds ratios. High-scoring cells were defined within each program as the top 5% by stGP score. Mann-Whitney $U$ tests compare the stGP scores of cells within each CN ($x_{\mathrm{CN}}$) against those outside that CN ($x_{\mathrm{rest}}$), and the resulting $p$ values are adjusted by the Benjamini-Hochberg procedure. Asterisks denote significant enrichment (*, adj. $P < 0.05$; **, adj. $P < 0.01$; ***, adj. $P < 0.001$). **g,h,** Distribution of high-scoring immune cells across CNs in the left (g) and right (h) kidney fits. Dot color and size denote the fraction of cells in each CN that were high scoring for the corresponding program. Asterisks denote one-sided Fisher exact tests for enrichment versus all other CNs, adjusted by the Benjamini-Hochberg procedure. The sham-inclusive analysis recovers the day-2 immune repair-transition program and its CN associations, supporting robustness to baseline inclusion.

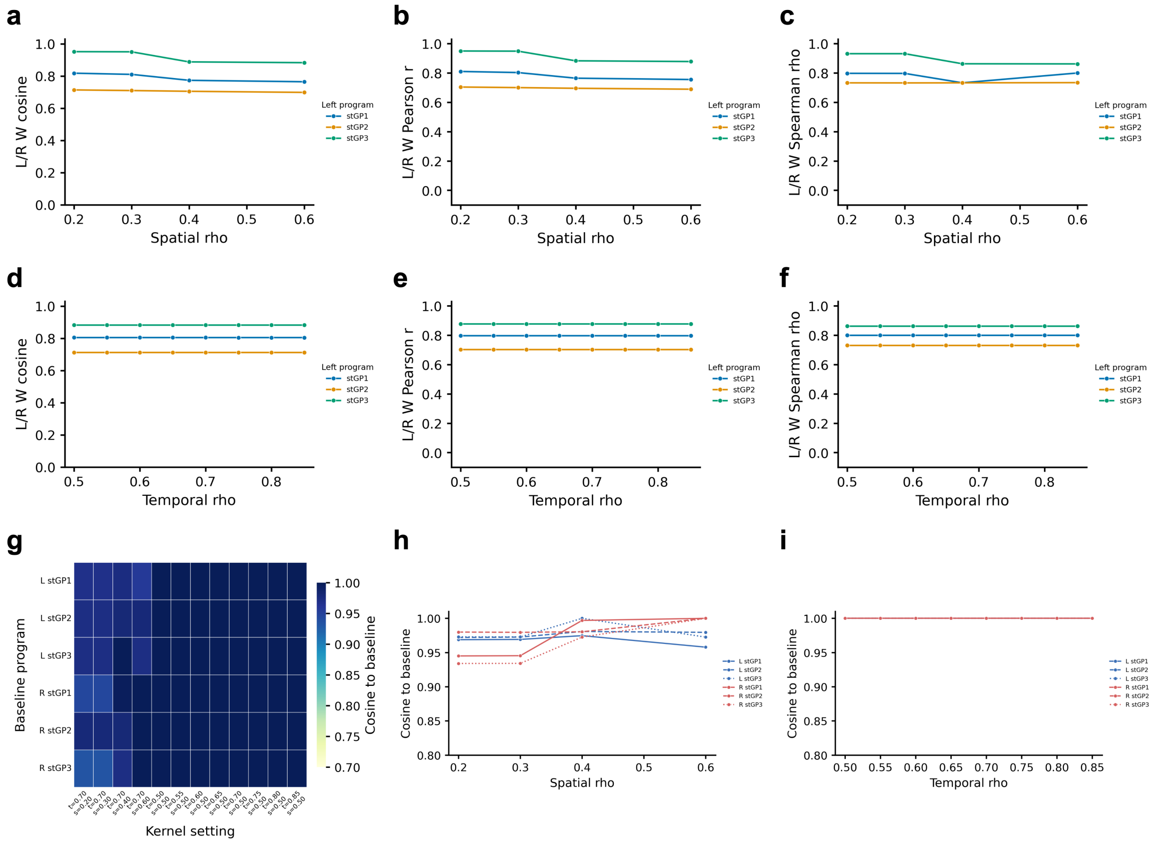

Supplementary Figure S21: **Immune program recovery is robust to spatial and temporal kernel settings.** stGP was refitted to left- and right-kidney immune cells across 12 temporal/spatial kernel settings. The baseline setting used temporal $\rho = 0.70$ and spatial $\gamma_{spa} = 0.50$. **a-c,** Left-right similarity of matched immune-program gene-loading vectors across spatial correlation settings, measured by (a) cosine similarity, (b) Pearson correlation, and (c) Spearman correlation. **d-f,** Left-right similarity across temporal correlation settings, measured by the same three metrics. **g,** Cosine similarity of gene-loading vectors between each tested kernel setting and the baseline setting. **h,i,** Program-wise cosine similarity to the baseline fit across spatial $\gamma_{spa}$ values (h) and temporal $\rho$ values (i). The high left-right similarity and high baseline similarity across settings indicate that the recovered immune programs are not driven by a narrow kernel choice.

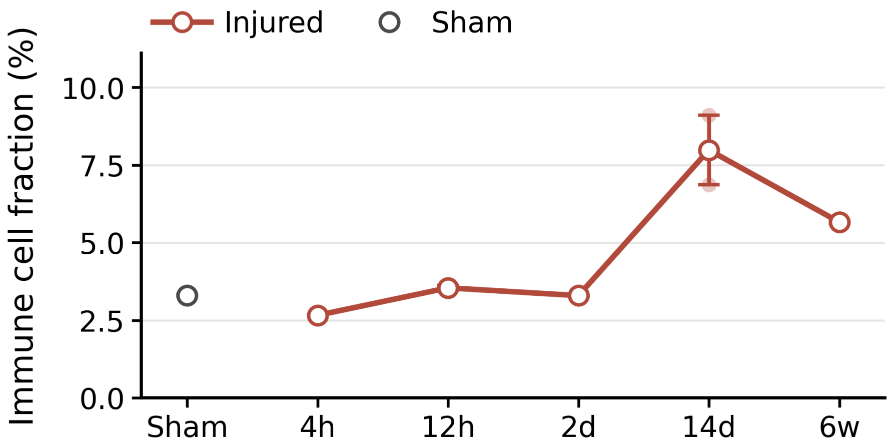

Supplementary Figure S22: **Immune-cell abundance peaks at day 14.** Immune-cell fraction was computed for each kidney section as the number of cells annotated as immune divided by the total number of cells in that section. The gray point denotes sham, and red open points denote injured kidneys at 4 h, 12 h, 2 days, 14 days and 6 weeks after BIRI. Error bars show the range across left and right kidneys. Immune-cell abundance increases most strongly at day 14 and remains elevated at 6 weeks.

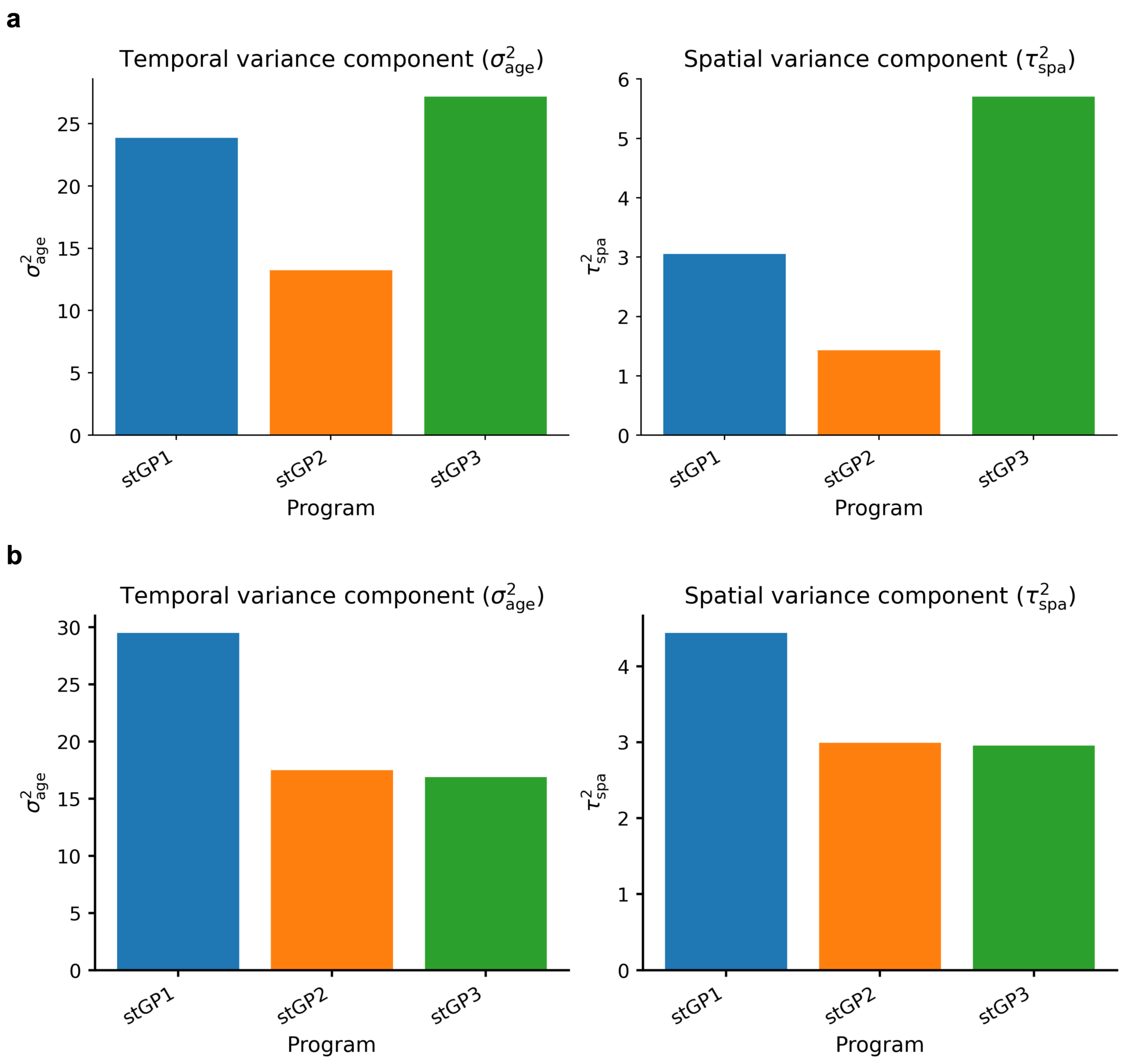

Supplementary Figure S23: **Estimated temporal and spatial variance components support stage-specific and spatially localized immune programs. a,b,** Estimated temporal variance component $\sigma^2_{\text{age}}$ and spatial variance component $\tau^2_{\text{spa}}$ for the three immune stGP programs in the left (a) and right (b) kidney fits. Larger $\sigma^2_{\text{age}}$ values indicate a stronger contribution from the shared post-injury temporal trajectory, whereas larger $\tau^2_{\text{spa}}$ values indicate stronger within-section spatial activities. All three programs show temporal and spatial contributions, with similar signal-to-noise ratios across the bilateral kidney fits.

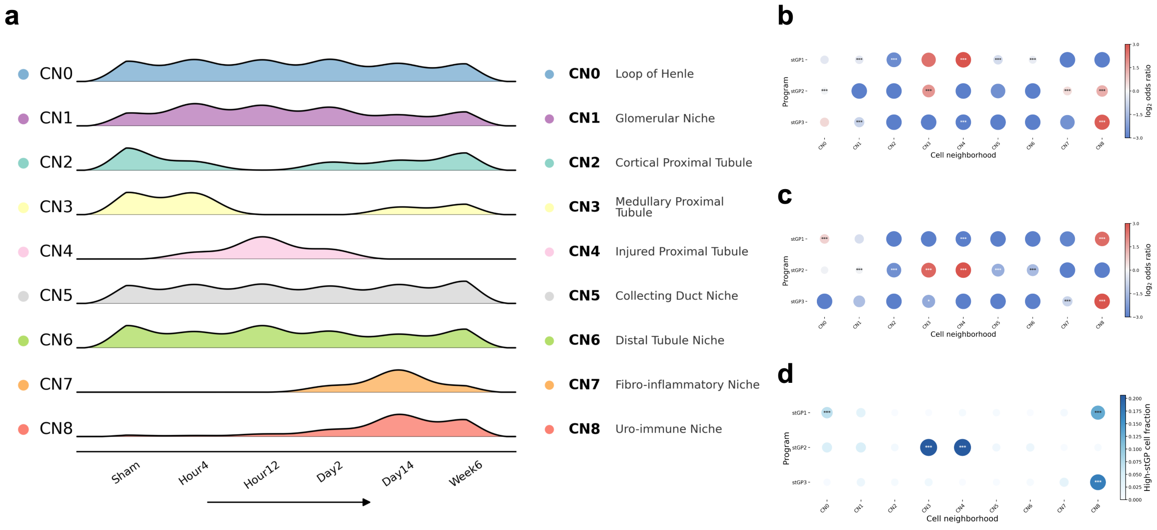

Supplementary Figure S24: **Cellular-neighborhood annotations define the tissue context of immune stGP programs. a,** Temporal distributions of the nine cellular neighborhoods (CNs) as well as their annotations in the kidney time course. The ridgeline summarizes CN abundance across sham, 4 h, 12 h, day 2, day 14 and week 6 stages. **b,c,** CN enrichment of high-scoring immune cells in the left (b) and right (c) kidney immune stGP fits, shown as $\log_2$ odds ratios. High-scoring cells were defined within each program as the top 5% by stGP score. Red indicates enrichment and blue indicates depletion. Mann-Whitney $U$ tests compare the stGP scores of cells within each CN ($x_{\mathrm{CN}}$) against those outside that CN ($x_{\mathrm{rest}}$), and the resulting $p$ values are adjusted by the Benjamini-Hochberg procedure. Asterisks denote significant enrichment (*, adj. $P < 0.05$; **, adj. $P < 0.01$; ***, adj. $P < 0.001$). **d,** Distribution of high-scoring immune cells across CNs for each matched immune program. Dot color and size denote the fraction of cells in each CN that were high scoring for the corresponding program. Asterisks denote one-sided Fisher exact tests for enrichment versus all other CNs, adjusted by the Benjamini-Hochberg procedure. Acute immune programs are enriched near medullary and injured proximal-tubule neighborhoods, whereas late remodeling programs are enriched in fibro-inflammatory and uro-immune neighborhoods.

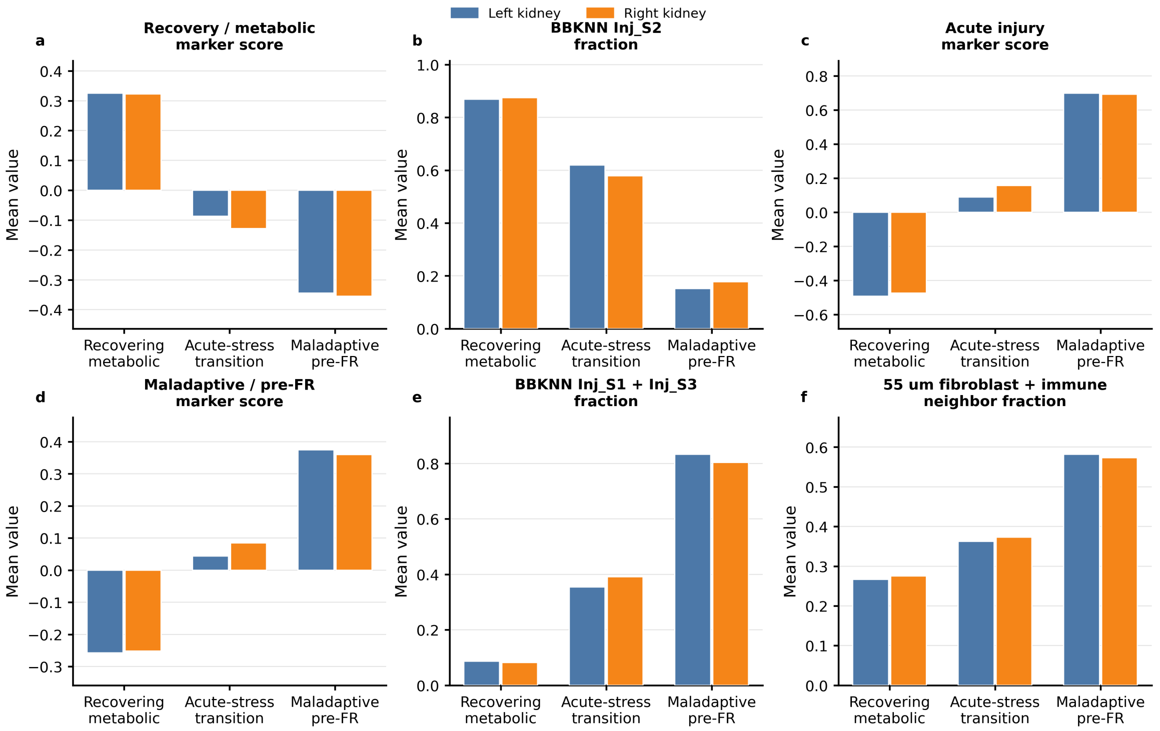

Supplementary Figure S25: **stGP spatial domains resolve a reproducible day-2 injured proximal tubule repair-state architecture across bilateral kidneys.** Day-2 injured proximal tubule (Inj-PT) cells in each side kidney were clustered into three domains in the stGP spatial embedding by $k$-nearest-neighbor spectral clustering. Because numeric domain IDs are replicate-specific, domains were relabeled as recovering metabolic, acute-stress transition or maladaptive pre-failed-repair (pre-FR) states using marker-program scores, BBKNN-derived reference proximal-tubule state labels and local tissue context. These validation features were not used to form the spectral clusters. **a,** Mean recovery/metabolic marker score, computed from z-scored expression of *Cxcl12*, *Haao*, *Kynu* and *Hmgcs2*. **b,** Fraction of cells assigned to the BBKNN Inj_S2 reference state. **c,** Mean acute-injury marker score, computed from z-scored expression of *Havcr1*, *Krt20* and *Plin2*. **d,** Mean maladaptive/pre-FR marker score, computed from z-scored expression of *Vcam1*, *Serpine1*, *Cd44* and *Klf5*. **e,** Fraction of cells assigned to the BBKNN Inj_S1 plus Inj_S3 reference states. **f,** Mean fraction of fibroblast and immune cells among reference neighbors within 55 $\mu$m of each Inj-PT cell. Blue and orange bars denote left and right kidneys, respectively. The recovering metabolic domain has the highest recovery score and Inj_S2 fraction, whereas the maladaptive pre-FR domain has the highest acute-injury and maladaptive scores, the largest Inj_S1 plus Inj_S3 fraction and the strongest fibroblast/immune proximity. The acute-stress transition domain shows intermediate features, supporting a replicated day-2 Inj-PT state architecture rather than side-specific clustering.

### D Supplementary Tables

Supplementary Table S1: **Runtime benchmark summary.**

| Data | Method | Runtime (min) | Note |
|---|---|---|---|
| Human brain – Excitatory neuron | stGP | 36.8 | – |
| | SpatialPCA | 8.8 | – |
| | MEFISTO | 53.0 | – |
| | Popari | 120.6 | – |
| | STAMP | 33.4 | – |
| Human brain – Oligodendrocyte | stGP | 3.6 | – |
| | SpatialPCA | 23.3 | – |
| | MEFISTO | 21.6 | – |
| | Popari | 41.4 | – |
| | STAMP | 30.7 | Invalid values |
| Mouse brain – Microglia | stGP | 40.5 | – |
| | SpatialPCA | 13.4 | – |
| | MEFISTO | 81.0 | – |
| | Popari | 48.4 | – |
| | STAMP | 61.8 | – |
